# Analysis of Plant-derived Phytochemicals as Anti-cancer Agents Targeting Cyclin Dependent Kinase-2, Human Topoisomerase IIa and Vascular Endothelial Growth Factor Receptor-2

**DOI:** 10.1101/2020.01.10.901660

**Authors:** Bishajit Sarkar, Md. Asad Ullah, Syed Sajidul Islam, Md. Hasanur Rahman

## Abstract

Cancer is caused by a variety of pathways, involving numerous types of enzymes, among them three enzymes: Cyclin dependent kinase-2 (CDK-2), Human topoisomerase IIα and Vascular Endothelial Growth Factor Receptor-2 (VEGFR-2) are three most common enzymes that are involved in the cancer development. Although many chemical drugs are available in the market, plant sources are known to contain a wide variety of agents that are known to possess anticancer activity. In this experiment, total thirty compounds were analysed against the mentioned enzymes using different tools of bioinformatics and *in silico* biology like molecular docking study, druglikeness property experiment, ADME/T test, PASS prediction and P450 site of metabolism prediction as well as DFT calculations to determine three best ligands that have the capability to inhibit the mentioned enzymes. Form the experiment, Epigallocatechin gallate was found to be the best ligand to inhibit CDK-2, Daidzein showed best inhibitory activities towards Human topoisomerase IIα and Quercetin was predicted to be the best agent against VEGFR-2. They were also predicted to be quite safe and effective agents to treat cancer. However, more *in vivo* and *in vitro* analysis are required to confirm their safety and efficacy in this regard.

## 1. Introduction

Cancer is defined as the uncontrolled proliferation and abnormal spread of the body’s specific cells. According to WHO, cancer was responsible for 13% of world deaths accounted in 2005. Moreover, projections have shown that cause-specific years of life lost (YLL) rate due to cancer would increase in 2005, 2015 and 2030. Millions of species of plants, animals, marine organisms and microorganisms act as attractive sources for new therapeutic candidate compounds. However, the development of novel agents from natural sources face many obstacles that are not usually met when one deals with synthetic compounds. Moreover, there may be difficulties with identification, isolation, assessing and obtaining the appropriate amounts of the active compound in the sample. (1, 2) The search for anti-cancer compounds from plant sources started in earnest in the 1950s with the discovery and development of the various natural compounds like vinca alkaloids, vinblastine, vincristine and cytotoxic podophyllotoxins. In the recent years, new technologies have been developed by the scientists to enhance natural product drug discovery in an industrial manner. Indeed, several new anticancer agents of natural origin have been introduced to the market recently and there is a promising pipeline of natural products in cancer-related clinical trials (3, 4, 5, 6). Future advances in the directed biosynthesis of small molecules will improve the ability of the scientists to control the shape and topology of various small molecules and thus creating new anti-cancer compounds that will interact specifically with biological targets. In the future, plants (300,000–500,000 such species) will continue to be a vital and valuable resource for anticancer drug discovery. More than 60 compounds from different plant sources are currently in the pipeline as potential anticancer agents (7, 8, 9, 10). Many chemical and synthetic drugs are already available for treating cancers i.e. alvocidib, lenvatinib and daunorubicin etc. These chemical drugs have many adverse effects like sepsis, diarrhea, stomach and bladder pain, hair loss, paralysis, joint pain etc. However, plant phytochemicals are considered as safe in this regard since they generally don’t possess any adverse effect to the human health in appropriate doses (11, 12, 13, 14). Therefore, using alternatives from plants can have great potential for cancer treatment. **Table 01** lists the potential phytochemicals used in the experiment.

**Table 01.**
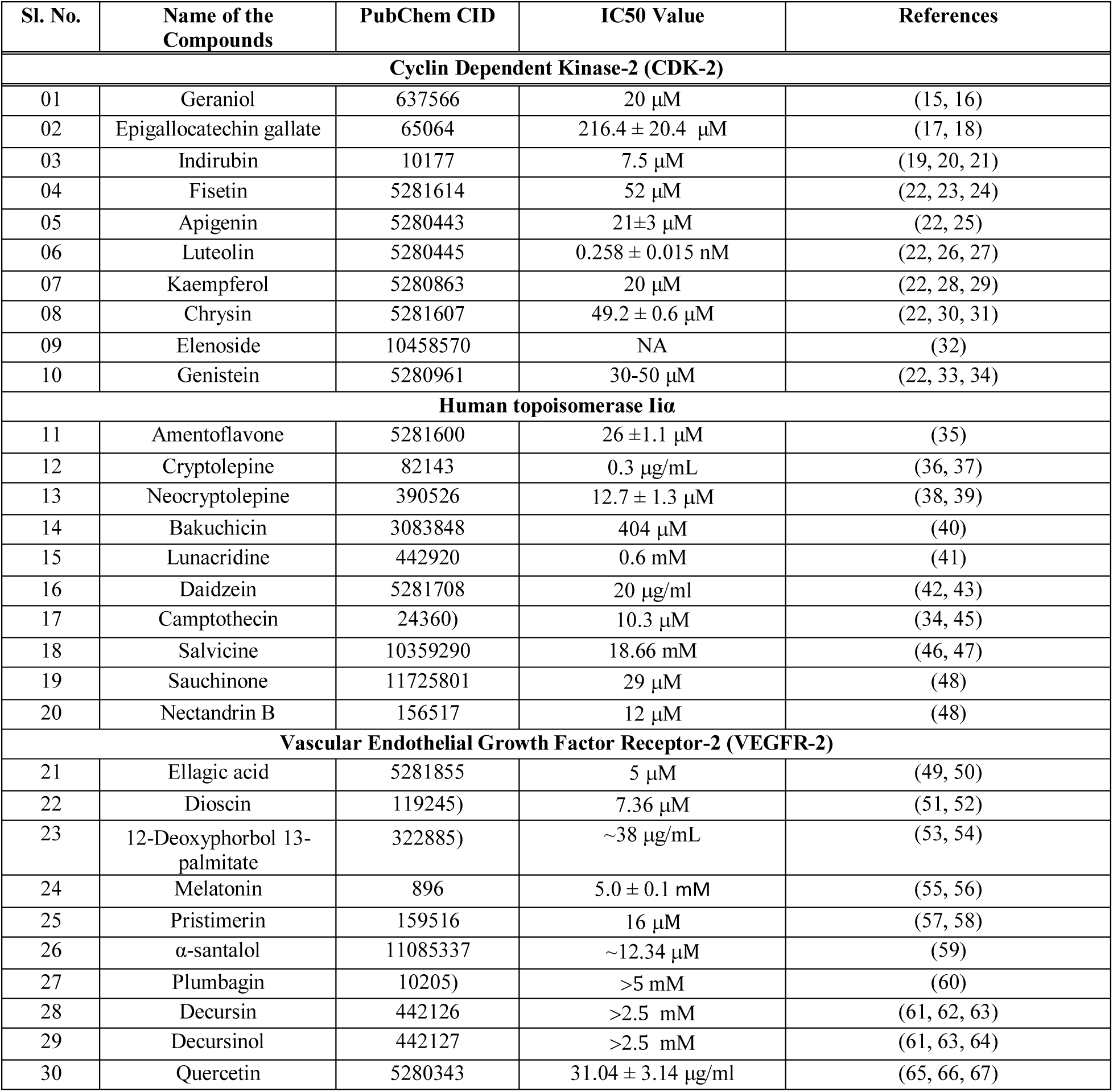
List of the plant derived anti-cancer agents that work via CDK-2, human topoisomerase IIa and VEGFR-2 pathways. NA; Not available.

| Sl. No. | Name of the Compounds | PubChem CID | IC50 Value | References |
| --- | --- | --- | --- | --- |
| **Cyclin Dependent Kinase-2 (CDK-2)** |  |  |  |  |
| 01 | Geraniol | 637566 | 20 μM | (15, 16) |
| 02 | Epigallocatechin gallate | 65064 | 216.4 ± 20.4 μM | (17, 18) |
| 03 | Indirubin | 10177 | 7.5 μM | (19, 20, 21) |
| 04 | Fisetin | 5281614 | 52 μM | (22, 23, 24) |
| 05 | Apigenin | 5280443 | 21±3 μM | (22, 25) |
| 06 | Luteolin | 5280445 | 0.258 ± 0.015 nM | (22, 26, 27) |
| 07 | Kaempferol | 5280863 | 20 μM | (22, 28, 29) |
| 08 | Chrysin | 5281607 | 49.2 ± 0.6 μM | (22, 30, 31) |
| 09 | Elenoside | 10458570 | NA | (32) |
| 10 | Genistein | 5280961 | 30-50 μM | (22, 33, 34) |
| **Human topoisomerase IIa** |  |  |  |  |
| 11 | Amentoflavone | 5281600 | 26 ± 1.1 μM | (35) |
| 12 | Cryptolepine | 82143 | 0.3 μg/mL | (36, 37) |
| 13 | Neocryptolepine | 390526 | 12.7 ± 1.3 μM | (38, 39) |
| 14 | Bakuchicin | 3083848 | 404 μM | (40) |
| 15 | Lunacridine | 442920 | 0.6 mM | (41) |
| 16 | Daidzein | 5281708 | 20 μg/ml | (42, 43) |
| 17 | Camptothecin | 24360 | 10.3 μM | (34, 45) |
| 18 | Salvicine | 10359290 | 18.66 mM | (46, 47) |
| 19 | Sauchinone | 11725801 | 29 μM | (48) |
| 20 | Nectandrin B | 156517 | 12 μM | (48) |
| **Vascular Endothelial Growth Factor Receptor-2 (VEGFR-2)** |  |  |  |  |
| 21 | Ellagic acid | 5281855 | 5 μM | (49, 50) |
| 22 | Dioscin | 119245 | 7.36 μM | (51, 52) |
| 23 | 12-Deoxyphorbol 13-palmitate | 322885 | ~38 μg/mL | (53, 54) |
| 24 | Melatonin | 896 | 5.0 ± 0.1 mM | (55, 56) |
| 25 | Pristimerin | 159516 | 16 μM | (57, 58) |
| 26 | α-santalol | 11085337 | ~12.34 μM | (59) |
| 27 | Plumbagin | 10205 | >5 mM | (60) |
| 28 | Decursin | 442126 | >2.5 mM | (61, 62, 63) |
| 29 | Decursinol | 442127 | >2.5 mM | (61, 63, 64) |
| 30 | Quercetin | 5280343 | 31.04 ± 3.14 μg/ml | (65, 66, 67) |

### 1.1. Role of Cyclin Dependent Kinase-2 (CDK-2) in cyclin/CDK Pathway and Its Involvement in Cancer

Cyclin/CDK pathway is one of the major cell cycle regulatory pathways, involving the cyclin-dependent kinases (CDKs), Retinoblastoma (Rb) tumor suppressor family and a family of transcription factors known as E2F. All these components of the pathway are essential for the passage of cells through the G1 to the S phase of the cell cycle. The CDK proteins are serine/threonine kinase that phosphorylate and thus inactivate the Rb protein. In the resting state of cell, Rb inhibits the activity of E2F protein forming a complex with it. The cyclin proteins can be of D type (cylcin D) and E type (cyclin E). Upon activation by the growth promoting signals or several mitogens, the cyclin D is found to form complex with CKD-4 and CKD-6. However, the cyclin E is found to be associated with CKD-2, when it is activated by active E2F. The cyclin D-CDK-4/6 and cyclin E-CDK-2 complexes phosphorylate and thereby inactivate the Rb protein. This inactivation causes the release of bound E2F transcription factor from the Rb protein. The released E2F later takes part in cell cycle progression. Moreover, E2F also promotes the activation of Cyclin E-CDK-2 complex, which in turn phosphorylates Rb protein and activate E2F transcription factor by feedback loop. Many inhibitors of the CDK proteins also takes part to regulate the cell cycle properly. The inhibitors repress the CDK proteins when there is no need for the cells to divide (68, 69, 70, 71, 72, 73, 74). The inhibitors are proteins from inhibitors of CDK-4 (INK4) and cyclin-dependent kinase inhibitor (CKI) families. CDK-4/6 is inhibited by p15/16 inhibitors and CDK-2 is inhibited by p21/p27 inhibitors. However, any type of mutation in the CDK genes causing hyperactivity or any type of mutation in the inhibitory genes, may lead to the uncontrolled proliferation of the cells, which can lead to different forms of cancers (75). For this reason, the targeting and inhibition of CDK-2 is a potential strategy for anticancer drug development (76) (**Figure 01**).

**Figure 01.**
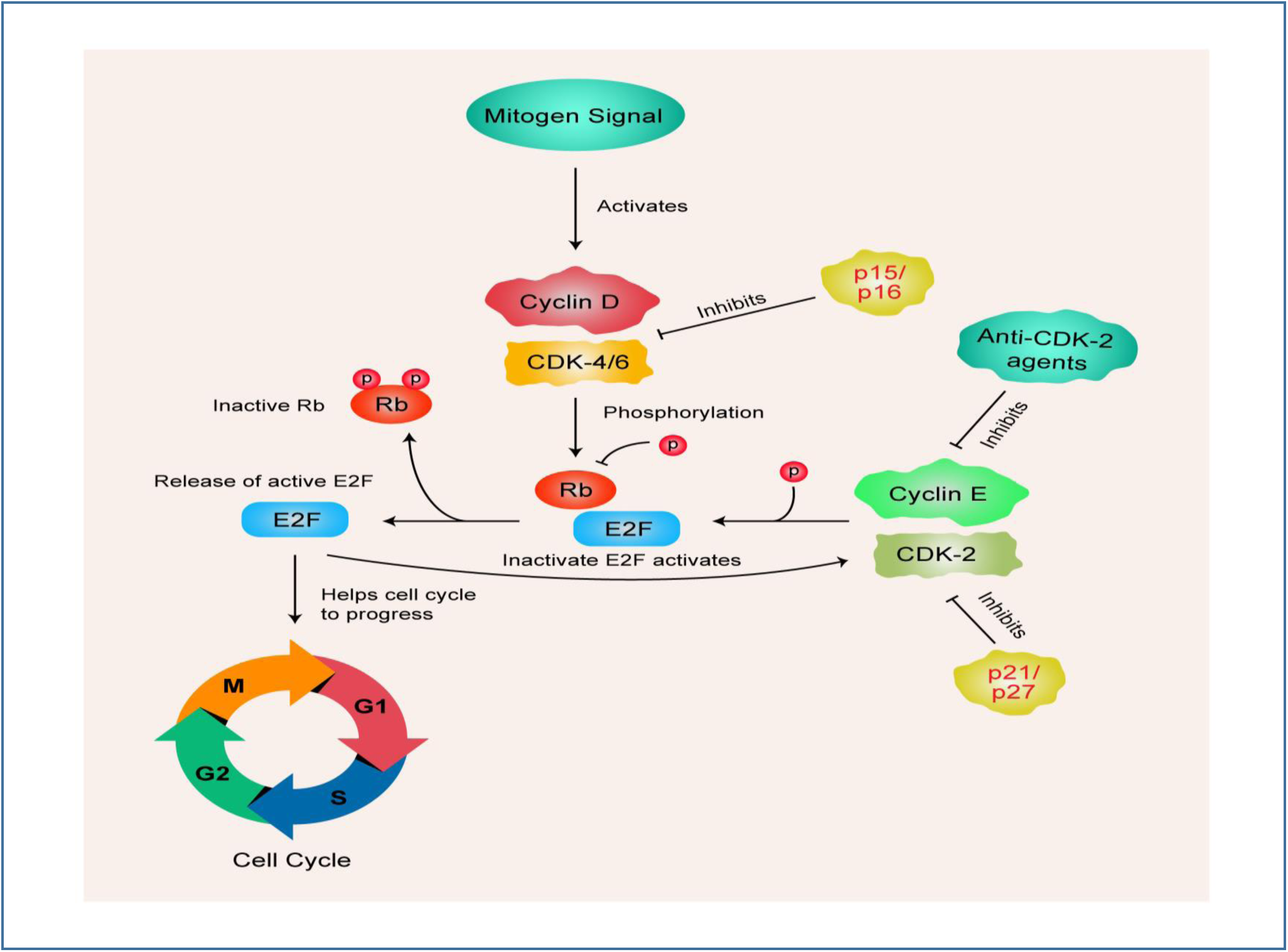
The cyclin/CDK signalling pathway. Upon activated by mitogen signal, the cyclin D-CDK-4/6 complex is activated and cause the inactivation of Rb by phosphorylation and thus release the active E2F, which takes part in cell cycle progression. However, E2F activates cyclin E-CDK-2 complex, which phosphorylates the Rb protein and activates the E2F in a feedback loop. P15/p16 inhibitors repress cyclin D-CDK-4/6 complex and p21/p27 inhibit cyclin E-CDK-2. Anti-CDK-2 agents inhibit the CDK-2 protein, thus can help in cancer treatment.

### 1.2. The DNA Topoisomerase IIα Pathway and Its Involvement in Cancer

Due to the supercoiled structure of the DNA molecules, it is necessary to unwind the double stranded DNA before replication, transcription, recombination and other processes. DNA topoisomerases are the enzymes that functions in unwinding, cutting, shuffling and relegating the DNA double helix structure. The human genome encodes six topoisomerases that are grouped into three types: type Iα, type Iβ and type IIα. DNA topoisomerase IIα is one the necessary topoisomerases that function in various cellular functions. However, it is a genotoxic enzyme which can lead to cancer development. When DNA topoisomerase II cuts the double stranded DNA, it may remain covalently attached to the broken end of the DNA. This reaction intermediate is known as the cleavage complex. If the amount of the cleavage complex in the cell falls too much, the cells are not able to divide into daughter cells due to mitotic failure, which results in the death of the cells. Moreover, if the amount of the cleavage complex increases too much, the temporary cleavage complex structures can become permanent double stranded breaks in the DNA. These double stranded breaks are caused by the faulty DNA tracking system which then initiate the faulty recombination and repair pathways of DNA, leading to cancer (**Figure 02**). For this reason, DNA topoisomerase IIα is a potential target for anti-cancer drug development (77, 78, 79, 80, 81).

**Figure 02.**
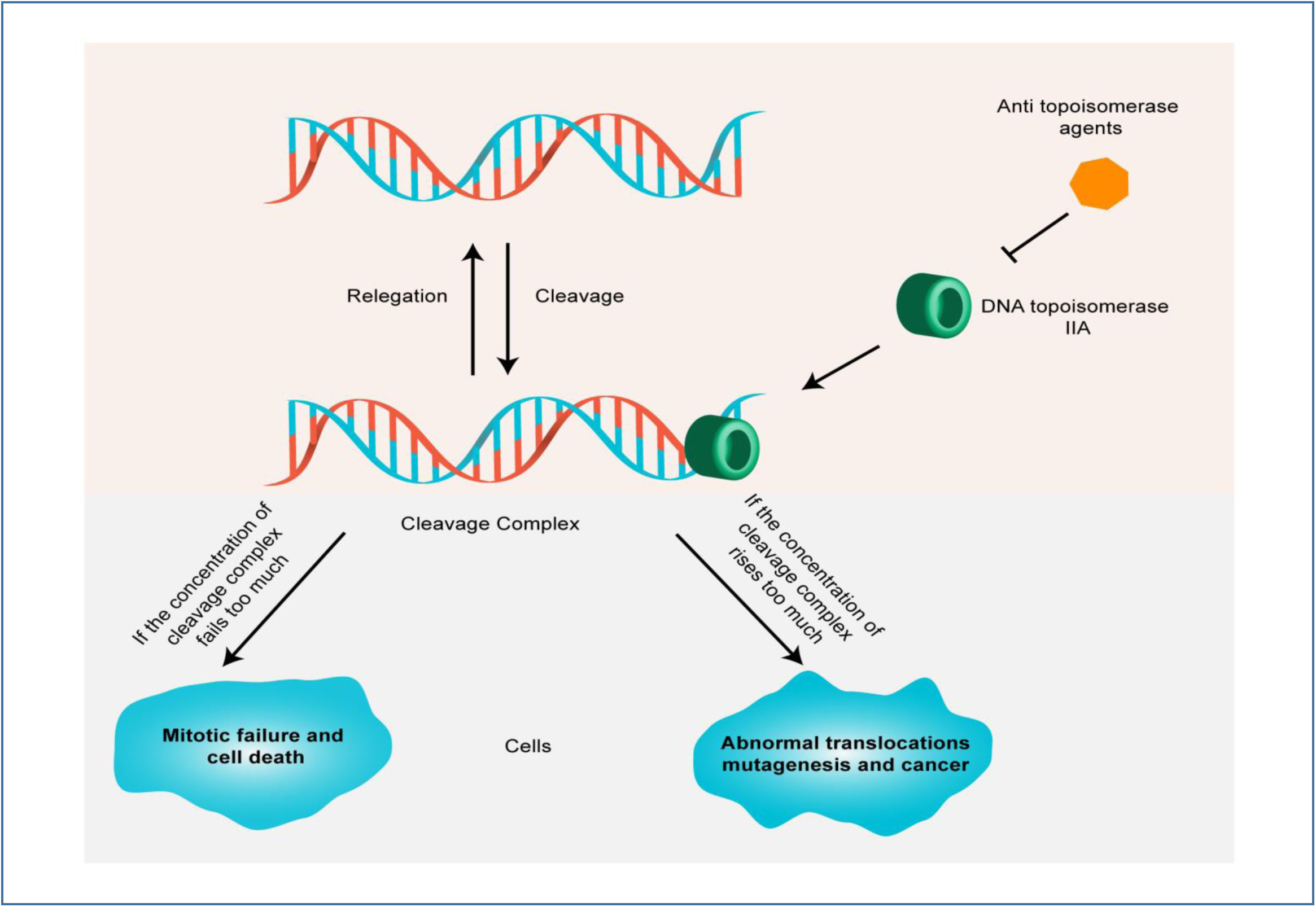
The DNA topoisomerase IIA pathway in cancer development. Upon the cleavage of the target DNA, the topoisomerase can remain bound to the cleaved ends of the DNA fragments and form cleavage complexes. If the concentration of cleavage complexes falls too much, then this may lead to cell death due to the mitotic failure. Moreover, if the concentration rises too much, abnormal translocations and mutagenesis may occur, which lead to cancer development. Anti-topoisomerase agents aid in cancer treatment by inhibiting the activity of DNA topoisomerase IIα.

### 1.3. The Role of Vascular Endothelial Growth Factor Receptor-2 (VEGFR-2) in Angiogenesis Pathway and Its Involvement in Cancer

Angiogenesis is the process of generating new capillary blood vessels (82). It plays important functions in organ development and differentiation during embryogenesis as well as wound healing and reproductive functions. However, angiogenesis is also responsible for a number of disorders including tumor formation, cancers, rheumatoid arthritis etc. Vascular Endothelial Growth Factor (VEGF) plays key role in angiogenesis process. VEGF protein has many isoforms and all of the isoforms mediate their effects by specific receptors known as VEGF receptors (VEGFRs). VEGFRs are receptor tyrosine kinases (RTKs) and there are three main isoforms: VEGFR-1, VEGFR-2, VEGFR-3. The expression of VEGF proteins are found to be dramatically increased in cancers like lung, thyroid, breast, ovary, kidney, uterine cancers etc. (83, 84). Since VEGF mediates its effects by binding to specific receptors (like VEGFR-2), inhibiting the actions of the receptors is thought to be a therapeutic target for cancer treatment (85). When VEGF protein binds with VEGFR-2, the VEGFR-2 becomes activated which then activates phosphatidylinositol 3-kinase (PI3K). PI3K further activates phosphoinositide-3-kinase (PIP3), which in turn activates the Akt/PKB (protein kinase B) signaling pathway. This pathway contributes to endothelial cell survival by activating proteins, like BAD (Bcl-2 associated death promoter) and caspase proteins. Moreover, the Akt/PKB signaling pathway can activate the endothelial nitric-oxide synthase (eNOS), which is responsible for vascular permeability. Both the endothelial cell survival and vascular permeability mechanisms contribute to the angiogenesis process. The binding of VEGF to VEGFR-2 can sometimes activate MAP kinase (mitogen activated protein kinase) pathway which is responsible for the proliferation of endothelial cells. In this pathway, activated VEGFR-2 activates phospholipase c-γ (PLC-γ). The PLC-γ then activates the protein kinase C (PKC). PKC further activates the proteins of MAP kinase pathway: RAF1, MEK, ERK, sequentially. This MAP kinase pathway causes the endothelial cell proliferation, which also contributes to the angiogenesis process (**Figure 03**) (86, 87, 88). Since VEGFR-2 is involved in angiogenesis process in cancer development, inhibition of VEGFR-2 is considered as therapeutic approach to treat cancer.

**Figure 03.**
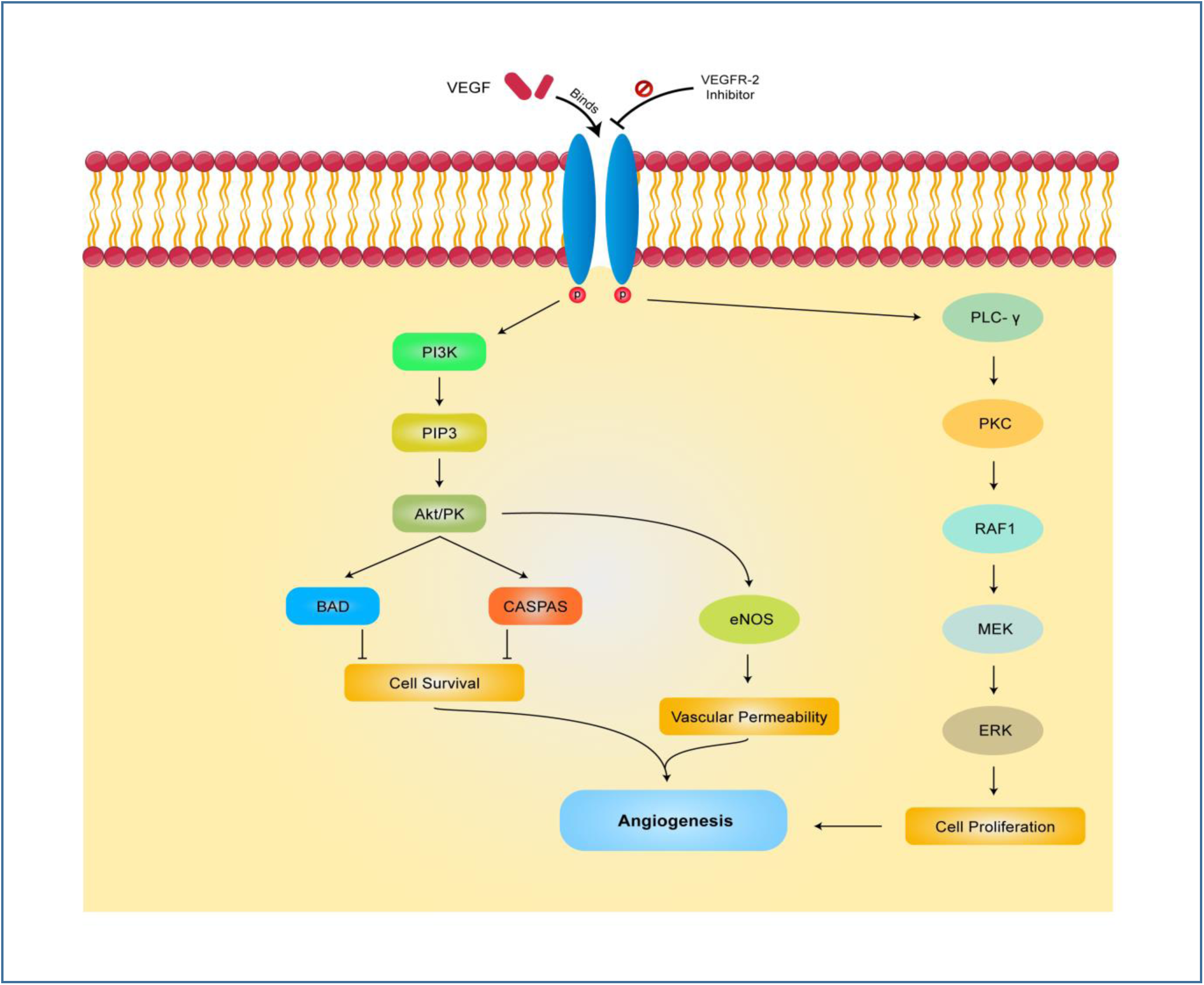
The angiogenesis pathway. The VEGF protein binds with VEGFR-2 and activates the receptor. The VEGFR-2 activates PI3K, which activates PIP3 and thus activating the Akt/PKB signaling pathway. This pathway contributes to endothelial cell survival by activating BAD and caspase proteins. Moreover, the Akt/PKB signaling pathway can activate eNOS, which is responsible for vascular permeability. Both the endothelial cell survival and vascular permeability mechanisms contribute to the angiogenesis process. Binding of VEGF to VEGFR-2 can sometimes activate MAP kinase pathway which is responsible for the proliferation of endothelial cells. The activated VEGFR-2 activates PLC-γ. The PLC-γ further activates PKC. PKC further activates RAF1, MEK, ERK, sequentially. This MAP kinase pathway causes the endothelial cell proliferation, which also contributes to the angiogenesis process. VEGFR-2 inhibitors inhibits VEGFR-2, thus aid in cancer treatment.

Three approved drugs were used as positive controls in this study: alvocidib (inhibits CDK-2), daunorubicin (inhibits human topoisomerase Iiα) and lenvatinib (inhibits VEGFR-2) (32, 89, 90).

## 2. Materials and methods

10 ligands (total) for each of the target molecule i.e., CDK-2, human topoisomerase IIa and VEGFR-2, were selected from literature that have already been proven to have inhibitory effects on the respective target molecule. Their IC50 values were collected by reviewing literatures discussing their anticancer potentiality. On sequential docking experiment one best ligand molecule was selected as the best inhibitor of respective target. Then their different drug like parameters were analysed in different experiments.

### 2.1. Protein Preparation and Ramachandran plot generation

Three dimensional structures of Cyclin-dependent kinase-2 (3EZV), Human topoisomerase II (1ZXM) and Vascular Endothelial Growth Factor Receptor-2 (2OH4) were downloaded (sequentially) in PDB format from protein data bank (www.rcsb.org). The proteins were then prepared and refined using the Protein Preparation Wizard in Maestro Schrödinger Suite 2018-4 (91). Bond orders were assigned and hydrogen molecules were added to heavy atoms as well as all the waters were deleted and the side chains were adjusted using Prime (92). Finally, the structure was optimized and then minimized using force field OPLS_2005. Minimization was done setting the maximum heavy atom RMSD (root-mean-square-deviation) to 30 Å and any remaining water less than 3 H-bonds to non-water was again deleted during the minimization step. After successful minimization, the proteins were used to generate Ramachandran plots for each of the protein by Maestro Schrödinger Suite 2018-4, keeping all the parameters as default.

### 2.2. Ligand Preparation

Three dimensional structures of 30 selected ligand molecules as well as controls were downloaded (sequentially) from PubChem database (www.pubchem.ncbi.nlm.nih.gov). These structures were then prepared using the LigPrep function of Maestro Schrödinger Suite (93). Minimized 3D structures of ligands were generated using Epik2.2 and within pH 7.0 +/-2.0 (94). Minimization was again carried out using OPLS_2005 force field which generated 32 possible stereoisomers.

### 2.3. Receptor Grid Generation

Grid usually confines the active site to shortened specific area of the receptor protein for the ligand to dock specifically. In Glide, a grid was generated using default Van der Waals radius scaling factor 1.0 and charge cutoff 0.25 which was then subjected to OPLS_2005 force field. A cubic box was generated around the active site (reference ligand active site). Then the grid box volume was adjusted to 15×15×15 for docking test.

### 2.4. Glide Standard Precision (SP) Ligand Docking, Prime MM-GBSA Calculation and Induced Fit Docking

SP adaptable glide docking was carried out using Glide in Maestro Schrödinger Suite (95). The Van der Waals radius scaling factor and charge cutoff were set to 0.80 and 0.15 respectively for all the ligand molecules. Final score was assigned according to the pose of docked ligand within the active site of the receptor.

This technique utilizes the docked complex and uses an implicit solvent which then assigns more accurate scoring function and improves the overall free binding affinity score upon the reprocessing of the complex. It combines OPLS molecular mechanics energies (E_MM_), surface generalized born solvation model for polar solvation (G_SGB_), and a nonpolar salvation term (G_NP_) for total free energy (ΔG_bind_) calculation. The total free energy of binding was calculated by the following equation: **ΔG_bind_ = G_complex_ – (G_protein_ – G_ligand_), where, G= E_MM_ + G_SGB_ + G_NP_** (96). Nine anticancer agents were selected on the basis of best MM-GBSA scores.

At this stage the docking parameters of our compounds under investigation was compared with 3 controls name with respective receptors.

To carry out the IFD of the nine selected ligand molecules, again OPLS_2005 force field was applied after generating grid around the co-crystallized ligand of the receptor and this time the best five ligands were docked rigidly. Receptor and Ligand Van Der Waals screening was set at 0.70 and 0.50 respectively, residues within 2 Å were refined to generate 2 best possible posses with extra precision. Best performing ligand was from each enzyme category was selected according to the IFD score and XP_Gscore_. The 3D representations of the best pose interactions between the ligands and their respective receptors were obtained using Discovery Studio Visualizer (97).

### 2.5. Ligand Based Drug Likeness Property and ADME/Toxicity Prediction

The drug likeness properties of the three selected ligand molecules were analyzed using SWISSADME server (http://www.swissadme.ch/) (98). The ADME/T for each of the ligand molecules was carried out using online based servers, admetSAR (http://lmmd.ecust.edu.cn/admetsar2/) and ADMETlab (http://admet.scbdd.com/) to predict their various pharmacokinetic and pharmacodynamic properties (99, 100). The absorption, distribution and metabolism properties were determined by both admetSAR server and excretion and toxicity properties were determined by ADMETlab server. The numeric and categorical values of the results given by ADMETlab server were changed into qualitative values according to the explanation and interpretation described in the ADMETlab server (http://admet.scbdd.com/home/interpretation/) for the convenience of interpretation.

### 2.6. PASS (Prediction of Activity Spectra for Substances) and P450 Site of Metabolism (SOM) prediction

The PASS (Prediction of Activity Spectra for Substances) prediction of the three best selected ligands were conducted by using PASS-Way2Drug server (http://www.pharmaexpert.ru/passonline/) by using canonical SMILES from PubChem server (https://pubchem.ncbi.nlm.nih.gov/) (101). To carry out PASS prediction, P_a_ (probability “to be active”) was kept greater than 70%, since the P_a_ > 70% threshold gives highly reliable prediction (102). In the PASS prediction study, both the possible biological activities and the possible adverse effects of the selected ligands were predicted. The P450 Site of Metabolism (SOM) of the three best selected ligand molecules were determined by online tool, RS-WebPredictor 1.0 (http://reccr.chem.rpi.edu/Software/RS-WebPredictor/) (103). The LD50 and Toxicity class was predicted using ProTox-II server (http://tox.charite.de/protox_II/) (104).

### 2.7. DFT Calculations

Minimized ligand structures obtained from LigPrep were used for DFT calculation using the Jaguar panel of Maestro Schrödinger Suite using Becke’s three-parameter exchange potential and Lee-Yang-Parr correlation functional (B3LYP) theory with 6-31G* basis set (105, 106, 107). Quantum chemical properties such as surface properties (MO, density, potential) and Multipole moments were calculated along with HOMO (Highest Occupied Molecular Orbital) and LUMO (Lowest Unoccupied Molecular Orbital) energy. Then the global frontier orbital was analyzed and hardness (**η**) and softness (**S**) of selected molecules were calculated using the following equation as per Parr and Pearson interpretation and Koopmans theorem (108, 109). The DFT calculation was done for the 3 best ligand molecules.

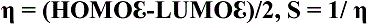

## 3. Results

### 3.1. **Ramachandran Plot and Molecular Docking Analysis**

After preparing the proteins, the Ramachandran plot for each of the receptor proteins was generated. In the plot, the orange regions represent “favored” regions, the yellow regions represent “allowed” regions and the white regions represent “disallowed” regions (110). CDK-2 protein generated Ramachandran plot with almost all of the amino acids in the “favored” region and no amino acids in the “disallowed region”. Human topoisomerase II generated Ramachandran plot with 15 amino acids in the “disallowed region”. It also had majority of the amino acids in the “favored” region. VEGFR-2 generated Ramachandran plot with only 4 amino acids in the “disallowed” region and most of the amino acids in the “favored” region (**Figure 04**).

**Figure 04.**
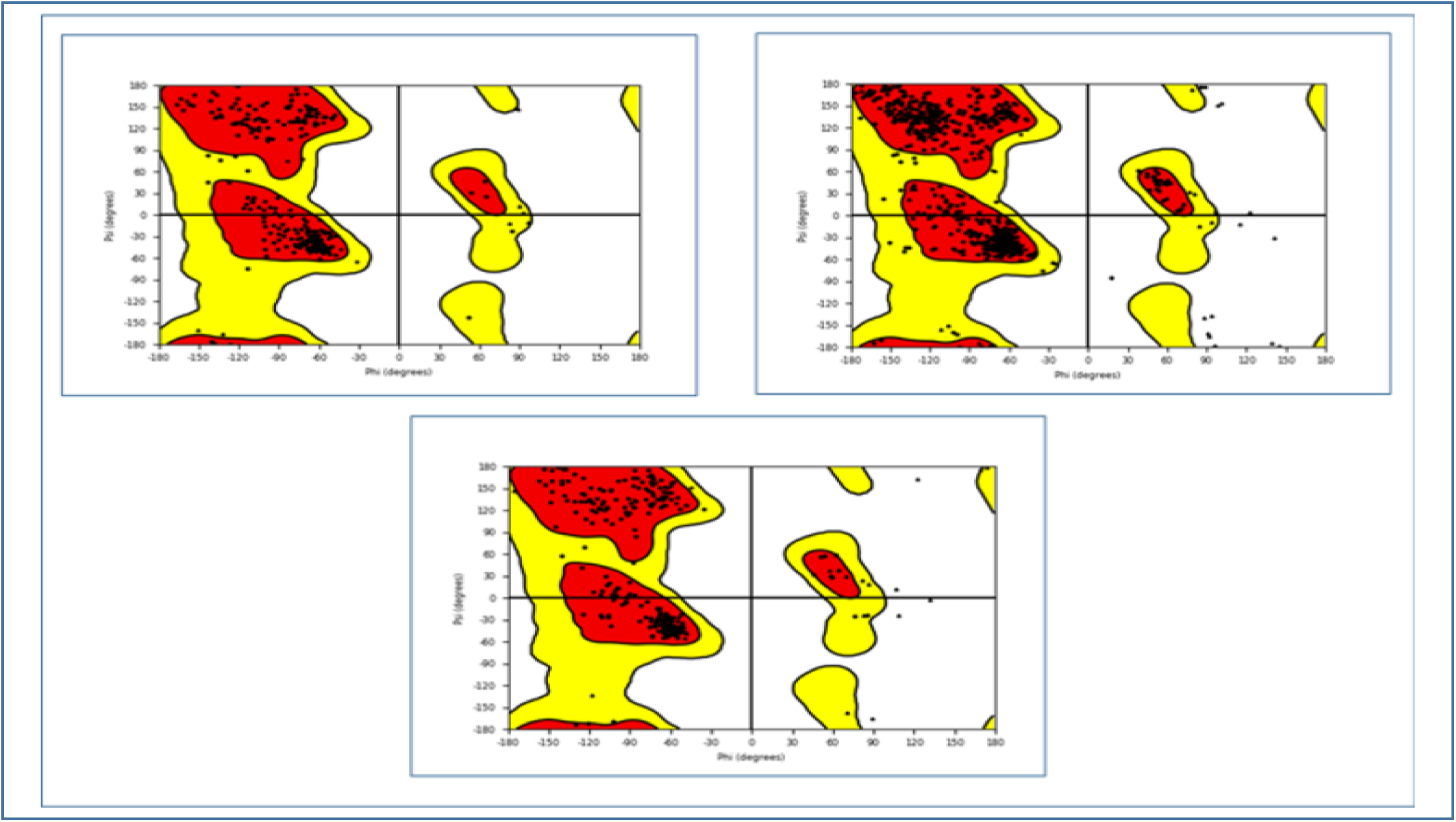
Ramachandran plot analysis of 1. CDK-2, 2. Human topoisomerase II, 3. VEGFR-2. Glycine and proline are represented as triangles and squares and all other amino acids are represented as spheres.

All the selected ligand molecules were docked successfully with their respective receptor proteins. The ligand molecules that had the lowest binding energy were considered the best ligand molecules in inhibiting their respective receptors since lower binding energy (docking score) corresponds to higher binding affinity (111). In the MM-GBSA study, the most negative ΔG_Bind_ score (the lowest score) is considered as the best ΔG_Bind_ score (112). IFD study is carried out to understand the accurate binding mode and to ensure the accuracy of active site geometry. The lowest values of IFD score and XP G_Score_ are considered as the best values (113, 114, 115, 116). Nine ligands: Geraniol, Epigallocatechin gallate and Indirubin (inhibit CDK-2), Daidzein, Camptothecin and Salvicine (inhibit human topoisomerase II) and Quercetin, Decursinol and Plumbagin (inhibit VEGFR-2), were initially selected based on the lower free binding energy and MM-GBSA study since they were reported to show comparable binding energy with respective controls (**Table 02**). Then these molecules were subjected to IFD study. Epigallocatechin gallate, Daidzein and Quercetin were considered as the three best ligand molecules from the IFD study among the nine initially selected ligands. The 3D representations as well as interaction of different amino acids with Epigallocatechin gallate, Daidzein and Quercetin are illustrated in **Figure 05**.

**Figure 05.**
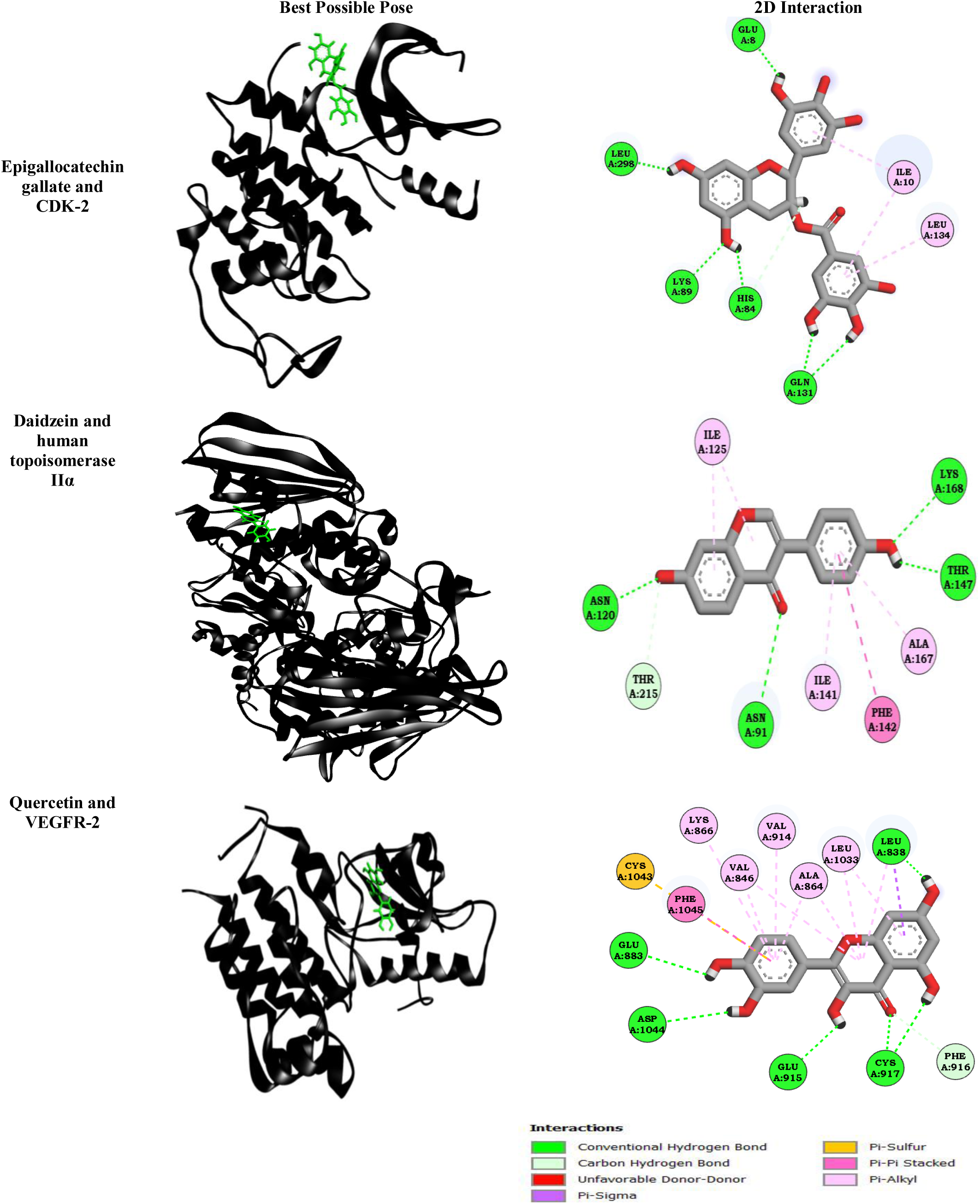
Best possible poses (left) and 2D interactions (right) between ligand and receptor molecules.

**Table 02.** The results of molecular docking study between the selected 30 ligands and their receptors.

| No | Name of ligand<br>(PubChem CID) | Name of receptor<br>(PDB ID) | Docking score/<br>binding energy<br>(Kcal/mol) | Glide energy<br>(Kcal/mol) | MM-GBSA<br>($\Delta G_{\text{Bind}}$ Score<br>Kcal/mol) |
| --- | --- | --- | --- | --- | --- |
| Control-1 | Alvocidib | CDK-2 (PDB ID: 3EZV) | -5.144 | -42.707 | -71.530 |
| 01 | Geraniol |  | -7.341 | -48.430 | -59.370 |
| 02 | Epigallocatechin gallate |  | -7.123 | -60.544 | -66.420 |
| 03 | Indirubin |  | -8.410 | -33.776 | -53.960 |
| 04 | Fisetin |  | -6.017 | -42.105 | -37.819 |
| 05 | Apigenin |  | -5.836 | -36.499 | -44.342 |
| 06 | Luteolin |  | -4.954 | -38.551 | -32.109 |
| 07 | Kaempferol |  | -4.046 | -39.418 | -38.334 |
| 08 | Chrysin |  | -5.893 | -34.446 | -46.700 |
| 09 | Elenoside |  | -7.036 | -46.435 | -35.451 |
| 10 | Genistein |  | -5.591 | -37.710 | -31.310 |
| Control-2 | Daunorubicin | Human topoisomerase II (PDB ID: 1ZXN) | -5.469 | -39.191 | -40.326 |
| 11 | Amentoflavone |  | -3.524 | -32.638 | -36.549 |
| 12 | Cryptolepine |  | -5.802 | -40.963 | -22.341 |
| 13 | Neocryptolepine |  | -4.977 | -39.058 | -37.330 |
| 14 | Bakuchicin |  | -5.638 | -37.756 | -40.004 |
| 15 | Lunacridine |  | -5.413 | -42.872 | -21.934 |
| 16 | Daidzein |  | -7.855 | -42.546 | -55.980 |
| 17 | Camptothecin |  | -7.630 | -48.500 | -40.223 |
| 18 | Salvicine |  | -6.969 | -42.072 | -44.550 |
| 19 | Sauchinone |  | -6.266 | -42.390 | -34.449 |
| 20 | Nectandrin B |  | -6.173 | -43.608 | -32.870 |
| Control-3 | Lenvatinib | VEGFR-2 (PDB ID: 2OH4) | -9.745 | -61.045 | -70.240 |
| 21 | Ellagic acid |  | -5.039 | -44.384 | -46.776 |
| 22 | Dioscin |  | -4.524 | -33.341 | -32.200 |
| 23 | 12-Deoxyphorbol 13-palmitate |  | -6.471 | -47.617 | -32.239 |
| 24 | Melatonin |  | -6.996 | -36.512 | -46.450 |
| 25 | Pristimerin |  | -6.179 | -37.520 | -33.984 |
| 26 | α-santalol | | -6.494 | -33.456 | -41.230 |
| 27 | Plumbagin |  | -7.848 | -40.639 | -48.910 |
| 28 | Decursin |  | -6.307 | -38.690 | -21.430 |
| 29 | Decursinol |  | -8.960 | -49.149 | -59.710 |
| 30 | Quercetin |  | -10.441 | -54.972 | -64.420 |

Now, these 3 best ligands (one from each of the receptor category) were used to in next phases of this experiment to analyze their drug potentials.

#### 3.1.1. Binding Mode of Best Ligands with Respective Targets

Epigallocatechin gallate docked with CDK-2 with an IFD score of −594.995 Kcal/mol and XP Gscore of −8.816 Kcal/mol. It formed six conventional hydrogen bonds with Lysine 89, Leucine 298, Histidine 84, Glutamic acid 08, Leucine 134 and Glutamine 131 (×2) at 1.82 Å, 1.53 Å, 2.13Å, 1.55 Å, 4.53 Å, 1.69 Å and 2.45 Å distance apart respectively within the binding pocket of CDK-2. Moreover, it also formed one non-conventional hydrogen bond with Histidine 84. Epigallocatechin gallate was also reported to form multiple hydrophobic interactions i.e., Pi-Alkyl with Isoleucine 10 and Leucine 34 amino acid residues within the binding cleft of CDK-2 (**Table 03**).

**Table 03.**
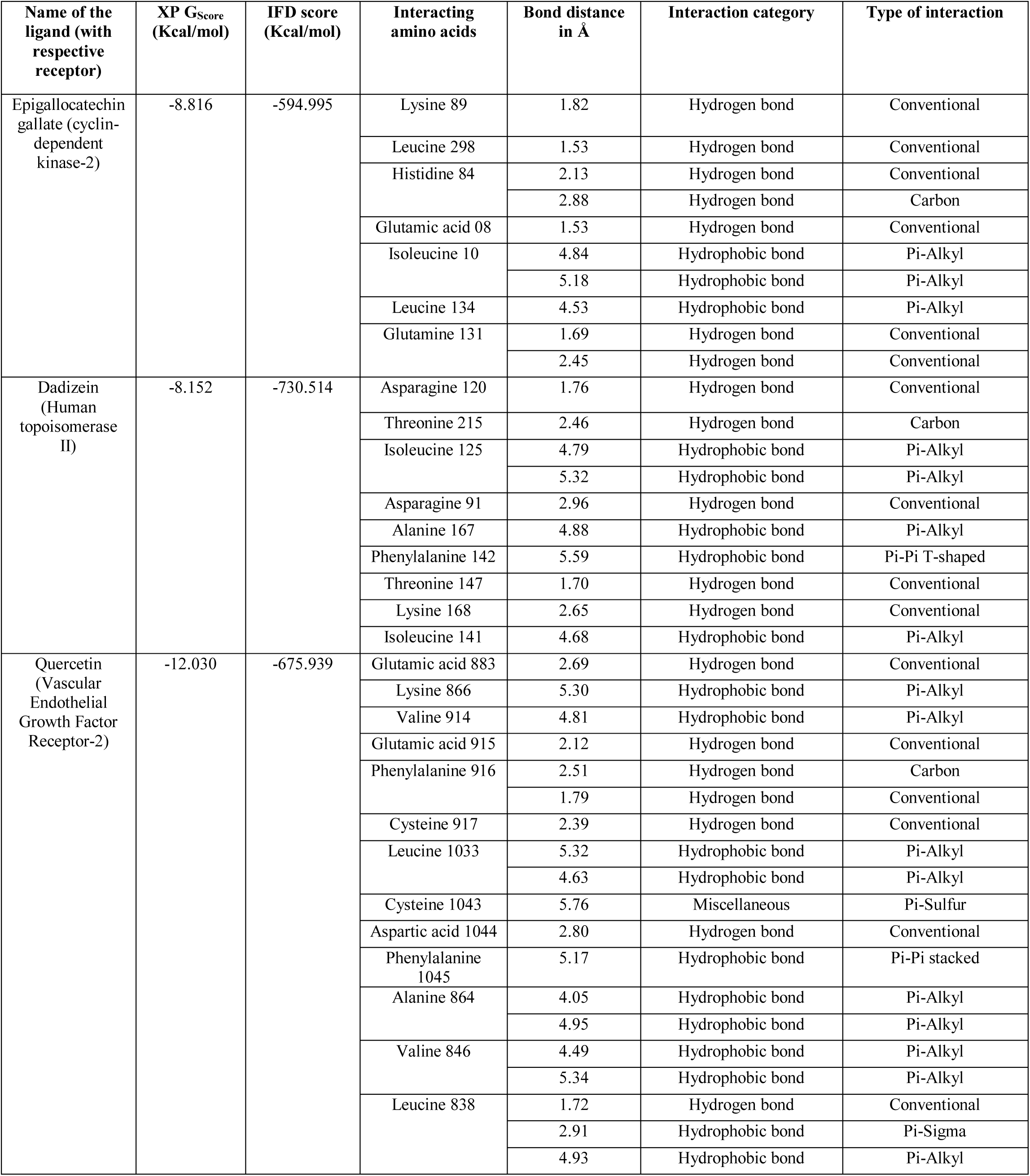
The results of docking studies between the three best ligands and their respective receptors, along with their interaction with different types of amino acids and the bonds formed between the ligands and the amino acids.

Daidzein docked with Topoisomerase IIa with an IFD score of – 730. 514 Kcal/mol and XP Gscore of −8.152 Kcal/mol. It formed four conventional hydrogen bonds with Asparagine 120, Asparagine 91, Threonine 147 and Lysine 168 at 1.82 Å, 1.76 Å, 2.96 Å, 1.71 Å and 2.65 Å distance apart respectively within the binding cleft of CDK-2. Daidzein was also reported to form multiple hydrophobic interactions i.e., Pi-Alkyl with Isoleucine 125 (×2) and Alanine 167 amino acid residues within the binding pocket of Human topoisomerase IIα (**Table 03**).

Quercetin docked with VEGFR-2 with an IFD score of −675.939 Kcal/ mol and XP Gscore of −12.030 Kcal/mol. It formed six conventional hydrogen bonds with Glutamic acid 883, Glutamic acid 915, Phenylalanine 916, Cysteine 1043, Aspartic acid 1044 and LEucine 838, at 2.69 Å, 2.12 Å, 1.79 Å, 5.76 Å, 2.80 Å and 1.72 Å distance apart respectively within the binding cleft of CDK-2. It also formed a non-conventional hydrogen bond with Phenylalanine 916 at 2.51 Å distance. It was also reported to form multiple hydrophobic interactions i.e., Pi-Alkyl with Leucine 838, Valine 914 and many other amino acid residues within the binding pocket of VEGFR-2 (**Table 03**).

### 3.2. Drug-likeness Properties

Among the three ligand molecules, only Epigallocatechin gallate violated the Lipinski’s rule of five (2 violations: number of hydrogen bond donors and acceptors). However, it showed the highest topological polar surface area (TPSA) value of 197.37 Å² (**Table 04**). Daidzein was shown to have highest LogP value and again Epigallocatechin Gallate showed highest molar refractivity. Daidzein and Quercetin each was reported to have 1 rotatable bond and Epigallocatechin gallate was predicted with 4 bonds.

**Table 04.** The drug-likeness properties of the best three ligands.

| Drug Likeness Properties | Rules | Epigallocatechin gallate | Daidzein | Quercetin |
| --- | --- | --- | --- | --- |
| Lipinski's rule of five | - | No (2 violations) | Yes | Yes |
| Molecular weight | <500 | 458.37 g/mol | 254.24 g/mol | 302.24 g/mol |
| Consensus Log $P_{o/w}$ | $\leq 5$ | 1.01 | 2.24 | 1.23 |
| Num. H-bond acceptors | <10 | 11 | 4 | 7 |
| Num. H-bond donors | <5 | 8 | 2 | 5 |
| No of rotatable bonds | $\leq 10$ | 4 | 1 | 1 |
| Molar Refractivity | 40-130 | 112.06 | 71.97 | 78.03 |
| TPSA ($\text{\AA}^2$) | - | 197.37 | 70.67 | 131.36 |

### 3.3 ADME/T Tests

The results of ADME/T test with probability scores are summarized in **Table 05**. In the absorption section, only daidzein showed positive Caco-2 permeability and all the three selected ligands were HIA positive. In the distribution section, all the molecules showed high capability to bind with plasma protein (PPB), however, all of them were not blood brain barrier permeable (BBB). In the metabolism section, only Epigallocatechin gallate was not inhibitory to CYP450 1A2 and quercetin was the only inhibitor of CYP450 3A4. None of the ligands were substrate for CYP450 2C9 and CYP450 2D6 and CYP450 2D6 had no predicted inhibitor. In the excretion section, Epigallocatechin gallate, Daidzein and Quercetin showed half-life of 1.7, 1.5 and 0.2 h, respectively. Only Epigallocatechin gallate showed hERG blocking capability, however, it didn’t show any human hepatotoxic activity (H-HT negative). Only daidzein showed negative result in the Ames mutagenicity test. However, all of them were DILI positive.

**Table 05.** The ADME/T test results of the best three ligand molecules.

| ADMET Class | Properties | Epigallocatechin gallate<br>(Probability Score) | Daidzein<br>(Probability Score) | Quercetin<br>(Probability Score) |
| --- | --- | --- | --- | --- |
| Absorption | Caco-2 permeability | Negative ( 0.9372) | Positive (0.931) | Negative (0.6417) |
|  | Pgp-inhibitor | Negative (0.5518) | Negative (0.916) | Negative (0.9191) |
|  | Pgp-substrate | Negative (0.8848) | Negative (0.936) | Negative (0.8360) |
|  | HIA (Human Intestinal Absorption) | Positive (0.9942) | Positive (0.989) | Positive (0.9833) |
| Distribution | PPB (Plasma Protein Binding) | High, 100% | High, 83.1% | High, 117.06% |
|  | BBB (Blood–Brain Barrier) | Negative (0.5464) | Negative (0.781) | Negative (0.463) |
| Metabolism | CYP450 1A2 inhibition | Negative (0.805) | Positive (0.910) | Positive (0.910) |
|  | CYP450 3A4 inhibition | Negative (0.766) | Negative (0.626) | Positive (0.695) |
|  | CYP450 3A4 substrate | Positive (0.567) | Negative (0.526) | Positive (0.556) |
|  | CYP450 2C9 inhibition | Negative (0.836) | Positive (0.975) | Negative (0.582) |
|  | CYP450 2C9 substrate | Negative (1.000) | Negative (0.820) | Negative (1.000) |
|  | CYP450 2C19 inhibition | Negative (0.808) | Positive (0.899) | Negative (0.902) |
|  | CYP450 2D6 inhibition | Negative (0.955) | Negative (0.919) | Negative (0.928) |
|  | CYP450 2D6 substrate | Negative (0.721) | Negative (0.800) | Negative (0.855) |
|  | Subcellular localization | Mitochondria | Mitochondria | Mitochondria |
| Excretion | T<sub>1/2</sub> (h) | 1.7 | 1.5 | 0.2 |
| Toxicity | hERG (hERG Blockers) | Blocker (0.812) | Non-blocker (0.394) | Non-blocker (0.371) |
|  | H-HT (Human Hepatotoxicity) | Negative (0.240) | Positive (0.784) | Positive (0.558) |
|  | Ames (Ames Mutagenicity) | Positive (0.560) | Negative (0.184) | Positive (0.740) |
|  | DILI (Drug Induced Liver Injury) | Positive (0.848) | Positive (0.948) | Positive (0.900) |

### 3.4. PASS and P450 Site of Metabolism Prediction

The predicted LD50 value for epigallocatechin gallate, daidzein and quercetin were 1000 mg/kg, 2430 mg/kg and 159 mg/kg, respectively. The prediction of activity spectra for substances (PASS prediction) was for the three selected ligands to identify 20 intended biological activities and 5 adverse & toxic effects. The PASS prediction results of all the three selected ligands are listed in **Table 06** and **Table 07**. The possible sites of metabolism by CYPs 1A2, 2A6, 2B6, 2C19, 2C8, 2C9, 2D6, 2E1 and 3A4 of Epigallocatechin gallate, Daidzein and Quercetin were determined (**Table 08**). The possible sites of metabolism by the isoforms are indicated by circles on the chemical structure of the molecule (117).

**Table 06.** The PASS prediction results showing the biological activities of the best three ligand molecules.

| Sl no | Biological activities | Epigallocatechin gallate |  | Daidzein |  | Quercetin |  |
| --- | --- | --- | --- | --- | --- | --- | --- |
|  |  | Predicted LD50: 1000 mg/kg |  | Predicted LD50: 2430 mg/kg |  | Predicted LD50: 159 mg/kg |  |
|  |  | Toxicity class: 4 |  | Toxicity class: 5 |  | Toxicity class: 3 |  |
|  |  | Pa | Pi | Pa | Pi | Pa | Pi |
| 01 | Antioxidant | 0.814 | 0.003 | - | - | 0.872 | 0.003 |
| 02 | Reductant | 0.944 | 0.002 | - | - | - | - |
| 03 | Anticarcinogenic | 0.841 | 0.004 | - | - | 0.757 | 0.007 |
| 04 | Antimutagenic | - | - | 0.836 | 0.003 | 0.940 | 0.001 |
| 05 | Chemopreventive | 0.860 | 0.003 | - | - | - | - |
| 06 | Membrane integrity agonist | 0.962 | 0.003 | 0.887 | 0.0014 | 0.973 | 0.002 |
| 07 | Hepatoprotectant | - | - | - | - | 0.706 | 0.007 |
| 08 | Mucomembranous protector | 0.950 | 0.003 | - | - | - | - |
| 09 | TP53 expression enhancer | 0.937 | 0.004 | 0.771 | 0.014 | 0.844 | 0.008 |
| 10 | Lipid peroxidase inhibitor | 0.946 | 0.002 | - | - | - | - |
| 11 | HIF1A expression inhibitor | - | - | - | - | 0.969 | 0.002 |
| 12 | APOA1 expression enhancer | 0.856 | 0.003 | - | - | - | - |
| 13 | Free radical scavenger | 0.934 | 0.001 | - | - | - | - |
| 14 | Peroxidase inhibitor | 0.759 | 0.007 | - | - | 0.962 | 0.001 |
| 15 | Oxidoreductase inhibitor | - | - | - | - | - | - |
| 16 | CYP1A1 inhibitor | - | - | 0.756 | 0.002 | 0.909 | 0.001 |
| 17 | CYP1A2 inhibitor | - | - | - | - | 0.909 | 0.003 |
| 18 | Antiviral | 0.771 | 0.003 | - | - | - | - |
| 19 | Cardioprotectant | - | - | - | - | 0.833 | 0.003 |
| 20 | Vasoprotector | - | - | 0.721 | 0.009 | 0.824 | 0.004 |

**Table 07.**
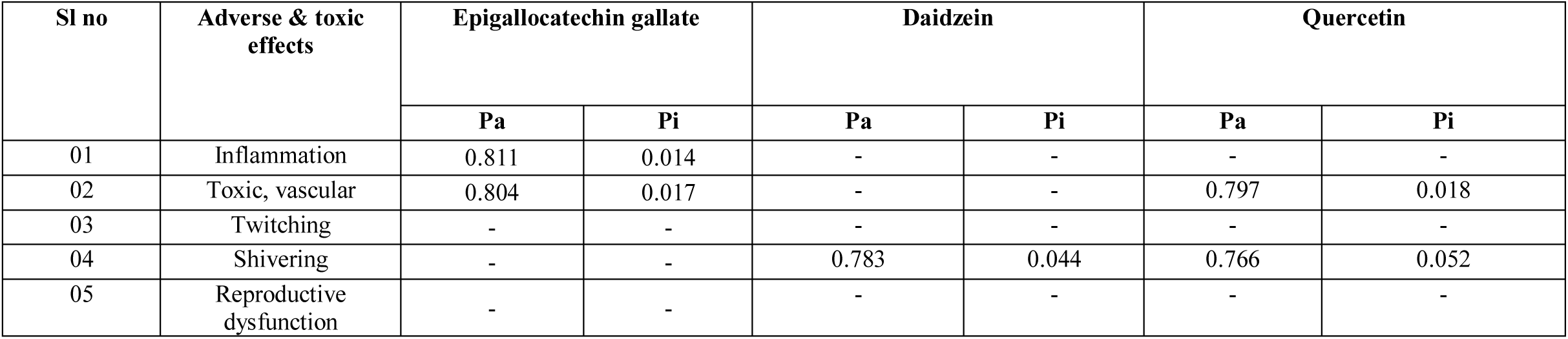
The PASS prediction results showing the adverse and toxic effects of the best three ligand molecules.

**Table 08.**
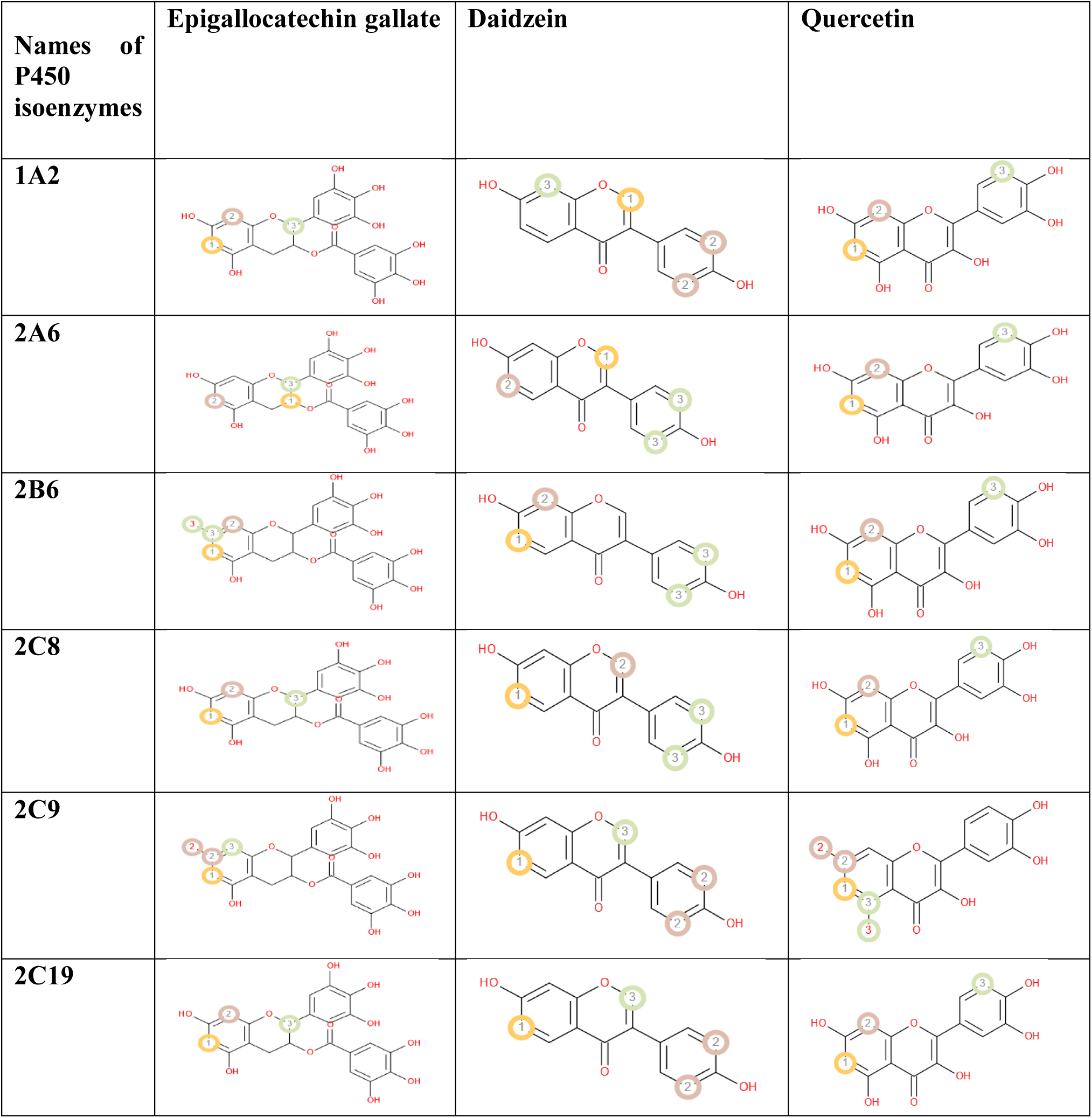
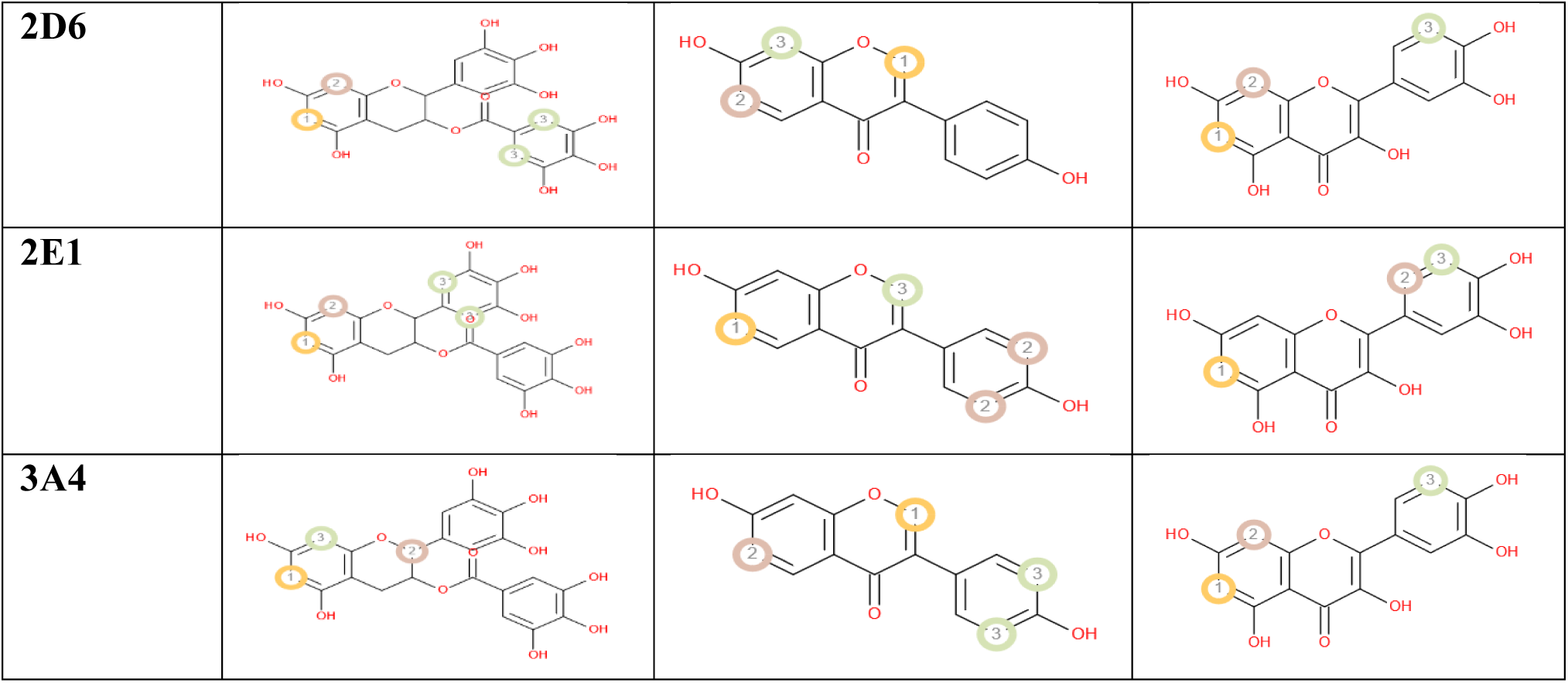
The P450 site of metabolism prediction of the best three ligand molecules.

### 3.6. Analysis of Frontier’s Orbitals

In the analysis of Frontier’s orbitals, the DFT calculations and HOMO-LUMO studies were conducted. The results of the DFT calculations are listed in **Table 09**. In these studies, Epigallocatechin gallate showed the lowest gap energy of 0.070 eV as well as the lowest dipole moment of 1.840 debye. On the other hand, quercetin generated the highest gap energy of 0.167 eV and the highest dipole moment of 5.289 debye. The order of gap energies and dipole moments of these three compounds were, epigallocatechin gallate < daidzein < quercetin (**Figure 06**).

**Figure 06.**
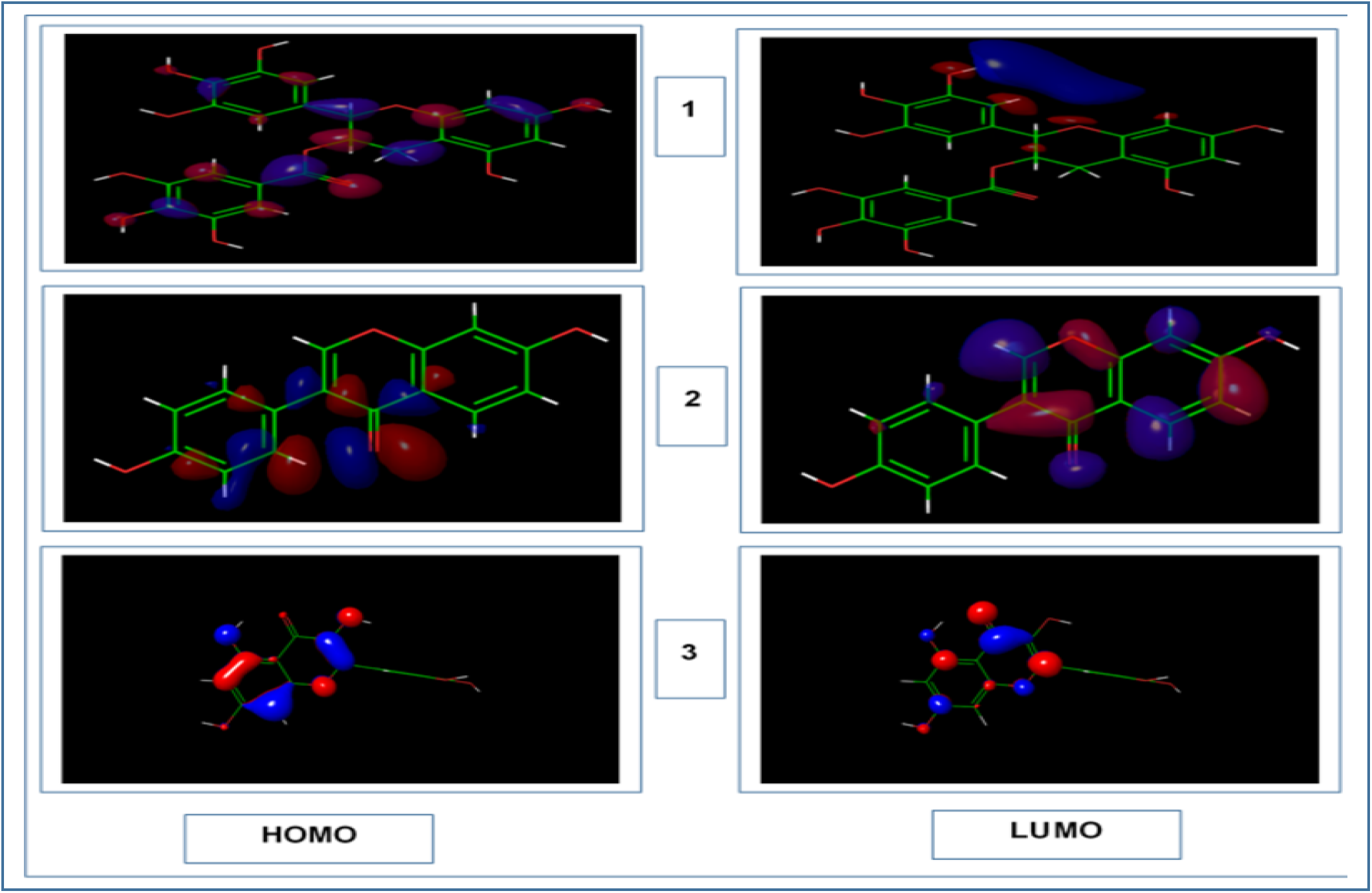
The HOMO and LUMO occupation; 1. Epigallocatechin gallate, 2. Daidzein and 3. Quercetin.

**Table 09.**
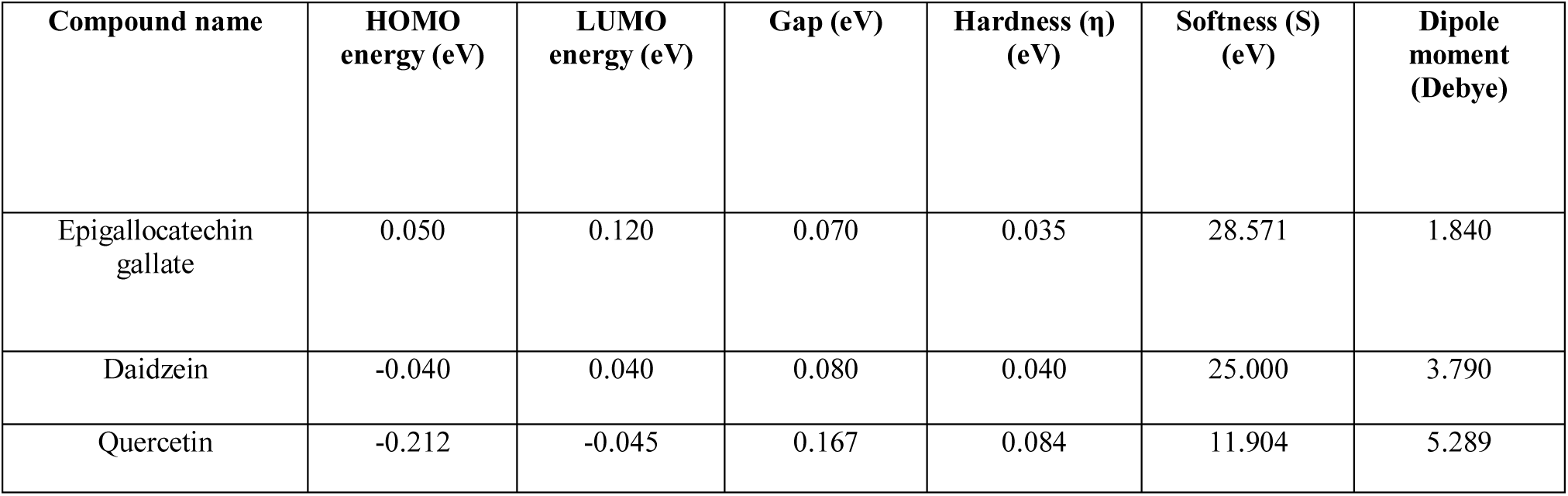
The results of the DFT calculations of the selected best three ligands.

## 4. Discussion

Molecular docking is an effective strategy in computer aided drug designing which works on specific algorithm and assigns affinity score depending on the poses od ligand inside the binding pocket of a target. Lowest docking score reflects highest affinity meaning that the complex remains more time in contact (118, 119).

In this study a total of 30 ligands targeting three macromolecules involved in cancer development were screened with the aid of molecular docking which generated comparable docking score as with positive controls (**Table 02**). At the initial step their quality was exemplified with the help of Ramachandran plot where they were predicted to perform well. Primarily, three ligands were selected for each receptor which were then subjected to IFD. Finally, Epigallocatechin gallate, Daidzein and Quercetin were selected as the best inhibitors of CDK-2, Human topoisomerase Iiα and VEGFR-2, respectively. Hydrogen and hydrophobic interactions are important for strengthening the receptor-ligand interactions (120). Selected best three ligands along with total ligands were predicted to form multiple hydrogen and hydrophobic interactions with the target molecules (**Table 02** and **Table 03**).

Estimation of the drug likeness properties facilitates the drug discovery and development process. Drug permeability through the biological barrier is influenced by the molecular weight and TPSA. The higher the molecular weight and TPSA, the lower the permeability of the drug molecule is and vice versa. Lipophilicity (expressed as LogP) affects the absorption of the drug molecule in the body and higher LogP associates with lower absorption. The number of hydrogen bond donors and acceptors beyond the acceptable range also affects the capability of a drug molecule to cross the cell membrane. The number of rotatable bonds also affects the druglikeness properties and the acceptable range is less than 10. Moreover, the Lipinski’s rule of five demonstrates that a successful drug molecule should have properties within the acceptable range of the five Lipinski’s rules (121, 122; Sarkar et al., 2019). Daidzein and Quercetin were reported to obey standard rule, whereas, Epigallocatechin gallate was reported to violate the rule which might subject it to further modification (**Table 04**).

The main purpose of conducting ADME/T tests is to determine the pharmacological and pharmacodynamic properties of a candidate drug molecule within a biological system. Therefore, it is a crucial determinant of the success of a drug discovery expenditure. BBB is the most important factor for those drugs that target primarily the brain cells. P-glycoprotein in the cell membrane aids in transporting many drugs, therefore, its inhibition affects the drug transport. The permeability of Caco-2 cell line indicates that the drug is easily absorbed in the intestine. Orally absorbed drugs travel through the blood circulation and deposit back to liver and are degraded by group of enzymes of Cytochrome P450 family and excreted as bile or urine. Therefore, inhibition of any of these enzymes of these enzymes affects the biodegradation of the drug molecule (123, 124). Moreover, if a compound is found to be substrate for one or more CYP450 enzyme or enzymes, then that compound is metabolized by the respective CYP450 enzyme or enzymes (125). A drug’s proficiency and pharmacodynamics are depended on the degree of its binding with the plasma protein. A drug can cross the cell layers or diffuse easily if it binds to the plasma proteins less efficiently and vice versa (126). Human intestinal absorption (HIA) is a crucial process for the orally administered drugs (127, 128, 129). Moreover, the half-life of a drug describes that the greater the half-life, the longer it would stay in the body and the greater its potentiality (130, 131, 132). HERG is a K^+^ channel found in the heart muscle and blocking the hERG signaling can lead to the cardiac arrhythmia (133, 134). Human hepatotoxicity (H-HT) involves any type of injury to the liver that may lead to organ failure and death (135, 136). Ames test is a mutagenicity assay that is used to detect the potential mutagenic chemicals (137). Drug induced liver injury (DILI) is the injury to the liver that are caused by administration of drugs. DILI is one of the causes that causes the acute liver failure (138). The results of ADME/T test are listed in **Table 05**. All of the three ligands were predicted to perform similar and sound in the ADME/T test.

Prediction of Activity Spectra for Substances or PASS prediction is a process that is used to estimate the possible profile of biological activities associated with drug-like molecules. Two parameters are used for the PASS prediction: Pa and Pi. The Pa is the probability of a compound “to be active” and Pi is the probability of a compound “to be inactive” and their values can range from zero to one (101). If the value of Pa is greater than 0.7, then the probability of exhibiting the activity of a substance in an experiment is higher (139). PASS was predicted for Epigallocatechin gallate, Daidzein and Quercetin. Both Epigallocatechin gallate and Quercetin showed similar performances with 12 biological activities and 2 toxic effects (**Table 06** and **Table 07**).

ProTox-II server estimates the toxicity of a chemical compound and classifies the compound into a toxicity class ranging from 1 to 6. The server classifies the compound according to the Globally Harmonized System of Classification and Labelling of Chemicals (GHS). According to the Globally Harmonized System of Classification and Labelling of Chemicals (GHS), since Epigallocactechin gallate had predicted toxicity class was of 4, it would be harmful if swallowed. With the predicted toxicity class of 5, Daidzein might be harmful if swallowed. And Quercetin, with its predicted toxicity class was 3, it was predicted to be toxic if swallowed (104, 140).

The Cytochrome P450 (Cyp450) is a superfamily of enzymes that comprises of 57 isoforms of P450 enzymes. These enzymes are heme-containing enzymes. They catalyze the phase-I metabolism of almost 90% of the marketed drugs and convert the lipophilic drugs to more polar compounds. Among the 57 isoforms, nine most prevalent isoforms are: CYPs 1A2, 2A6, 2B6, 2C19, 2C8, 2C9, 2D6, 2E1 and 3A4 (141, 142). All three best selected ligands showed multiple SOMs for these nine isoforms of P450, which indicates that they might be metabolized well by the body.

Frontier orbitals study or DFT calculation is an essential method of determining the pharmacological properties of various small molecules. HOMO and LUMO help to study and understand the chemical reactivity and kinetic stability of small molecules. The term ‘HOMO’ directs to the regions on a small molecule which may donate electrons during a complex formation and the term ‘LUMO’ indicates the regions on a small molecule that may receive electrons from the electron donor(s). The difference in HOMO and LUMO energy is known as gap energy that corresponds to the electronic excitation energy. The compound that has the greater orbital gap energy, tends to be energetically unfavourable to undergo a chemical reaction and vice versa (107, 143, 144, 145, 146, 148). All of the ligands were reported to have significant energy gap indicating their possibility to undergo a chemical reaction (**Table 09** and **Figure 06**).

Finally, all the best performed ligands were analyzed in different post-screening study and they’re predicted to perform sound. Overall, this study recommends Epigallocatechin gallate, Daidzein and Quercetin as the best inhibitors of CDK-2, Human topoisomerase Iiα and VEGFR-2, respectively among all selected ligands which could be potential natural plant-derived compounds to treat cancer. However, other compounds could also be investigated as they were also showed convincing docking scores (**Table 02**). Further in vivo and in vitro experiments might be required to strengthen the findings of this study.

## 5. Conclusion

In the experiment, 30 anti-cancer agents were selected to analyse against three enzymes, CDK-2, human topoisomerase IIa and VEGFR-2, of three different pathways that lead to cancer development. 10 ligands were studied for each of the enzyme group using different approaches used in computer-aided drug designing. Upon continuous computational experimentation, Epigallocatechin gallate, Daidzein and Quercetin were predicted to the best inhibitors of CDK-2, Human topoisomerase IIα and VEGFR-2 respectively. Then their drug potentiality was checked in different post-screening studies where they were also predicted to perform quite similar and sound. However, the authors suggest more *in vivo* and *in vitro* researches to be performed on these agents as well as the other remaining agents to finally confirm their potentiality, safety and efficacy.

## Acknowledgements

Authors are thankful to Swift Integrity Computational Lab, Dhaka, Bangladesh, a virtual platform of young researchers, for providing the tools.

## 6. Conflict of interest

Bishajit Sarkar declares that he has no conflict of interest. Md. Asad Ullah declares that he has no conflict of interest. Syed Sajidul Islam declares that he has no conflict of interest. Md. Hasanur Rahman declares that he has no conflict of interest.

## References

1. World health statistics 2006. 2006, 1–80, WHO (http://www.who.int/)

2. Da Rocha., A.B., Lopes, R.M. and Schwartsmann, G. Natural products in anticancer therapy. Curr. Opin. Pharmacol. 2001, 1(4), 364–369.

3. Cragg, G., M.; Newman, D., J. Plants as a source of anti-cancer agents. J. Ethnopharmacol. 2005, 100(1-2), 72–79.

4. Pan, L.; Chai, H.; Kinghorn, A., D. The continuing search for antitumor agents from higher plants. Phytochem. Lett. 2010, 3(1), 1–8.

5. Sarkar, B.; Ullah, M., A.; Islam, M., S.; Hossain, S; Nafi-Ur-Rahman, M. Anticancer potential of medicinal plants from Bangladesh and their effective compounds against cancer. J. Pharmacogn. Phytochem. 2019, 8(3), 827–833.

6. Pan, L.; Chai, H., B.; Kinghorn, A., D. Discovery of new anticancer agents from higher plants. Front. Biosci. (Scholar edition) 2012, 4, 142.

7. Fischbach, M., A.; Walsh, C., T. Directing biosynthesis. Science 2006, 314(5799), 603–605.

8. Fabricant, D., S.; Farnsworth, N., R. The value of plants used in traditional medicine for drug discovery. Environ. Health Persp. 2001, 109(suppl 1), 69–75.

9. Tan, G.; Gyllenhaal, C.; Soejarto, D., D. Biodiversity as a source of anticancer drugs. Curr. Drug Targets 2006, 7(3), 265–277.

10. Saklani, A.; Kutty, S., K. Plant-derived compounds in clinical trials. Drug Discov. Today 2008, 13(3-4), 161–171.

11. https://ascopost.com/News/58966. Accessed on: 3 March, 2020.

12. https://www.mayoclinic.org/drugs-supplements/lenvatinib-oral-route/side-effects/drg-7764?p=1 2013. Accessed on: 3 March, 2020.

13. https://www.drugs.com/sfx/daunorubicin-side-effects.html. Accessed on: 3 March, 2020.

14. Karimi A, Majlesi M, Rafieian-Kopaei M. Herbal versus synthetic drugs; beliefs and facts. Journal of nephropharmacology. 2015;4(1):27.

15. Qi F, Yan Q, Zheng Z, Liu J, Chen Y, Zhang G. Geraniol and geranyl acetate induce potent anticancer effects in colon cancer Colo-205 cells by inducing apoptosis, DNA damage and cell cycle arrest. J BUON. 2018 Mar 1;23(2):346–52.

16. Duncan RE, Lau D, El-Sohemy A, Archer MC. Geraniol and β-ionone inhibit proliferation, cell cycle progression, and cyclin-dependent kinase 2 activity in MCF-7 breast cancer cells independent of effects on HMG-CoA reductase activity. Biochemical pharmacology. 2004 Nov 1;68(9):1739–47.

17. Amin, A.; Gali-Muhtasib, H.; Ocker, M.; Schneider-Stock, R. Overview of major classes of plant-derived anticancer drugs. Int. J. Biomed. Sci. 2009, 5(1), 1.

18. Ramirez-Mares, M., V.; Chandra, S.; de-Mejia, E., G. In vitro chemopreventive activity of Camellia sinensis, Ilex paraguariensis and Ardisia compressa tea extracts and selected polyphenols. Mutat. Res. 2004, 554(1-2), 53–65.

19. Ponnusamy, K.; Petchiammal, C.; Mohankumar, R.; Hopper, W. In vitro antifungal activity of indirubin isolated from a South Indian ethnomedicinal plant Wrightia tinctoria R. Br. J. Ethnopharmacol. 2010, 132(1), 349–354.

20. Jautelat, R.; Brumby, T.; Schäfer, M.; Briem, H.; Eisenbrand, G.; Schwahn, S.; Krüger, M.; Lücking, U.; Prien, O.; Siemeister, G. From the insoluble dye indirubin towards highly active, soluble CDK2-inhibitors. ChemBioChem. 2005, 6(3), 531–540.

21. Hoessel R, Leclerc S, Endicott JA, Nobel ME, Lawrie A, Tunnah P, Leost M, Damiens E, Marie D, Marko D, Niederberger E. Indirubin, the active constituent of a Chinese antileukaemia medicine, inhibits cyclin-dependent kinases. Nature cell biology. 1999 May;1(1):60–7.

22. K-Jain, S.; B-Bharate, S.; a-Vishwakarma, R. Cyclin-dependent kinase inhibition by flavoalkaloids. Mini-rev. Med. Chem. 2012, 12(7), 632–649.

23. Lu, X.; Jung, J., I.; Cho, H., J.; Lim, D., Y.; Lee, H., S.; Chun, H., S.; Kwon, D., Y.; Park, J., H. Fisetin inhibits the activities of cyclin-dependent kinases leading to cell cycle arrest in HT-29 human colon cancer cells. J. Nutr. 2005, 135(12), 2884–2890.

24. Rengarajan T, Yaacob NS. The flavonoid fisetin as an anticancer agent targeting the growth signaling pathways. European journal of pharmacology. 2016 Oct 15;789:8–16.

25. Shukla, S.; Gupta, S.; Apigenin-induced cell cycle arrest is mediated by modulation of MAPK, PI3K-Akt, and loss of cyclin D1 associated retinoblastoma dephosphorylation in human prostate cancer cells. Cell cycle 2007, 6(9), 1102–1114.

26. Lin, Y.; Shi, R.; Wang, X.; Shen, H., M. Luteolin, a flavonoid with potential for cancer prevention and therapy. Curr. Cancer Drug Tar. 2008, 8(7), 634–646.

27. Saewan N, Koysomboon S, Chantrapromma K. Anti-tyrosinase and anti-cancer activities of flavonoids from Blumea balsamifera DC. J Med Plants Res. 2011 Mar 18;5(6):1018–25.

28. Cho, H., J.; Park, J., H., Y. Kaempferol induces cell cycle arrest in HT-29 human colon cancer cells. J. Cancer prev. 2013, 18(3), 257.

29. Yang J, Xiao P, Sun J, Guo L. Anticancer effects of kaempferol in A375 human malignant melanoma cells are mediated via induction of apoptosis, cell cycle arrest, inhibition of cell migration and downregulation of m-TOR/PI3K/AKT pathway. J BUON. 2018 Jan 1;23:218–23.

30. Weng, M., S.; Ho, Y., S.; Lin, J., K. Chrysin induces G1 phase cell cycle arrest in C6 glioma cells through inducing p21Waf1/Cip1 expression: involvement of p38 mitogen-activated protein kinase. Biochem. Pharmacol. 2005, 69(12), 1815–1827.

31. Samarghandian S, Azimi Nezhad M, Mohammadi G. Role of caspases, Bax and Bcl-2 in chrysin-induced apoptosis in the A549 human lung adenocarcinoma epithelial cells. Anti-Cancer Agents in Medicinal Chemistry (Formerly Current Medicinal Chemistry-Anti-Cancer Agents). 2014 Jul 1;14(6):901–9.

32. Khan, W.; Ashfaq, U., A.; Aslam, S.; Saif, S.; Aslam, T.; Tusleem, K.; Maryam, A.; ul-Qamar, M., T. Anticancer screening of medicinal plant phytochemicals against Cyclin-Dependent Kinase-2 (CDK2): An in-silico approach. *Adv*. Life Sci. 2017, 4(4), 113–119.

33. Choi, Y., H.; Lee, W., H.; Park, K., Y.; Zhang, L. p53-independent induction of p21 (WAF1/CIP1), reduction of cyclin B1 and G2/M arrest by the isoflavone genistein in human prostate carcinoma cells. Jpn. J. Cancer Res. 2000, 91(2), 164–173.

34. Sarkar FH, Adsule S, Padhye S, Kulkarni S, Li Y. The role of genistein and synthetic derivatives of isoflavone in cancer prevention and therapy. Mini reviews in medicinal chemistry. 2006 Apr 1;6(4):401–7.

35. Grynberg, N., F.; Carvalho, M., D.; Velandia, J., R.; Oliveira, M., C.; Moreira, I., C.; Braz-Filho, R.; Echevarria, A. DNA topoisomerase inhibitors: biflavonoids from Ouratea species. Braz. J. Med. Boil. Res. 2002, 35(7), 819–822.

36. Kirby, G., C.; Paine, A.; Warhurst, D., C.; Noamese, B., K.; Phillipson, J., D. In vitro and in vivo antimalarial activity of cryptolepine, a plant-derived indoloquinoline. Phytother. Res. 1995, 9(5), 359–363.

37. Bonjean, K.; De-Pauw-Gillet, M., C.; Defresne, M., P.; Colson, P.; Houssier, C.; Dassonneville, L.; Bailly, C.; Greimers, R.; Wright, C.; Quetin-Leclercq, J.; Tits, M. The DNA intercalating alkaloid cryptolepine interferes with topoisomerase II and inhibits primarily DNA synthesis in B16 melanoma cells. Biochemistry 1998, 37(15), 5136–5146.

38. Bailly, C.; Laine, W.; Baldeyrou, B.; De, M., P., G.; Colson, P.; Houssier, C.; Cimanga, K.; Van, S., M.; Vlietinck, A., J.; Pieters, L. DNA intercalation, topoisomerase II inhibition and cytotoxic activity of the plant alkaloid neocryptolepine. Anti-cancer Drug Des. 2000, 15(3), 191–201.

39. Dassonneville, L.; Lansiaux, A.; Wattelet, A.; Wattez, N.; Mahieu, C.; Van-Miert, S.; Pieters, L.; Bailly, C. Cytotoxicity and cell cycle effects of the plant alkaloids cryptolepine and neocryptolepine: relation to drug-induced apoptosis. Eur. J. Pharmacol. 2000, 409(1),9–18.

40. Sun, N., J.; Woo, S., H.; Cassady, J., M; Snapka, R., M.; DNA polymerase and topoisomerase II inhibitors from Psoralea corylifolia. J. Nat. Prod. 1998, 61(3), 362–366.

41. Prescott, T., A.; Sadler, I., H.; Kiapranis, R.; Maciver, S., K. Lunacridine from Lunasia amara is a DNA intercalating topoisomerase II inhibitor. J. Ethnopharmacol. 2007, 109(2), 289–294.

42. Jo, J., Y.; Gonzalez-de-Mejia, E.; Lila, M., A. Catalytic inhibition of human DNA topoisomerase II by interactions of grape cell culture polyphenols. J. Agr. Food Chem. 2006, 54(6), 2083–2087.

43. Vissac-Sabatier C, Bignon YJ, Bernard-Gallon DJ. Effects of the phytoestrogens genistein and daidzein on BRCA2 tumor suppressor gene expression in breast cell lines. Nutrition and cancer. 2003 Mar 1;45(2):247–55.

44. Sugimoto, Y.; Tsukahara, S.; Oh-hara, T.; Liu, L., F.; Tsuruo, T. Elevated expression of DNA topoisomerase II in camptothecin-resistant human tumor cell lines. Cancer Res. 1990, 50(24), 7962–7965.

45. Zheng MS, Lee YK, Li Y, Hwangbo K, Lee CS, Kim JR, Lee SK, Chang HW, Son JK. Inhibition of DNA topoisomerases I and II and cytotoxicity of compounds from Ulmus davidiana var. japonica. Archives of pharmacal research. 2010 Sep 1;33(9):1307–15.

46. Lu, H., R.; Meng, L., H.; Huang, M.; Zhu, H.; Miao, Z., H.; Ding, J. DNA damage, c-myc suppression and apoptosis induced by the novel topoisomerase II inhibitor, salvicine, in human breast cancer MCF-7 cells. Cancer Chemoth. Pharm. 2005, 55(3), 286–294.

47. Zhang, Y.; Wang, L.; Chen, Y.; Qing, C. Anti-angiogenic activity of salvicine. Pharm. Boil. 2013, 51(8), 1061–1065.

48. Lee, Y., K.; Seo, C., S.; Lee, C., S.; Lee, K., S.; Kang, S., J.; Jahng, Y.; Chang, H., W.; Son, J., K. Inhibition of DNA topoisomerases I and II and cytotoxicity by lignans from Saururus chinensis. Arch. Pharm. Res. 2009, 32(10), 1409.

49. Maas, J., L.; Galletta, G., J.; Stoner, G., D. Ellagic acid, an anticarcinogen in fruits, especially in strawberries: a review. HortScience. 1991, 26(1), 10–14.

50. Labrecque, L.; Lamy, S.; Chapus, A.; Mihoubi, S.; Durocher, Y.; Cass, B.; Bojanowski, M., W.; Gingras, D.; Béliveau, R. Combined inhibition of PDGF and VEGF receptors by ellagic acid, a dietary-derived phenolic compound. Carcinogenesis. 2005, 26(4), 821–826.

51. Tong, Q.; Qing, Y.; Wu, Y.; Hu, X.; Jiang, L.; Wu, X. Dioscin inhibits colon tumor growth and tumor angiogenesis through regulating VEGFR2 and AKT/MAPK signaling pathways. Toxicol. Appl. Pharm. 2014, 281(2), 166–173.

52. Cho, J.; Choi, H.; Lee, J.; Kim, M., S.; Sohn, H., Y.; Lee, D., G. The antifungal activity and membrane-disruptive action of dioscin extracted from Dioscorea nipponica. Biomembrane 2013, 1828(3), 1153–1158.

53. Xu, H., Y.; Pan, Y., M.; Chen, Z., W.; Lin, Y.; Wang, L., H.; Chen, Y., H.; Jie, T., T.; Lu, Y., Y. Liu, J., C. 12-Deoxyphorbol 13-palmitate inhibit VEGF-induced angiogenesis via suppression of VEGFR-2-signaling pathway. J. Ethnopharmacol. 2013, 146(3), 724–733.

54. Xu HY, Chen ZW, Li H, Zhou L, Liu F, Lv YY, Liu JC. 12-Deoxyphorbol 13-palmitate mediated cell growth inhibition, G2-M cell cycle arrest and apoptosis in BGC823 cells. European journal of pharmacology. 2013 Jan 30;700(1-3):13–22.

55. Cerezo, A.; Hornedo-Ortega, R.; Álvarez-Fernández, M.; Troncoso, A.; García-Parrilla, M. Inhibition of VEGF-Induced VEGFR-2 activation and HUVEC migration by melatonin and other bioactive indolic compounds. Nutrients 2017, 9(3), 249.

56. Plaimee P, Weerapreeyakul N, Barusrux S, Johns NP. Melatonin potentiates cisplatin-induced apoptosis and cell cycle arrest in human lung adenocarcinoma cells. Cell proliferation. 2015 Feb;48(1):67–77.

57. Mu X, Shi W, Sun L, Li H, Jiang Z, Zhang L. Pristimerin, a triterpenoid, inhibits tumor angiogenesis by targeting VEGFR2 activation. Molecules. 2012 Jun;17(6):6854–68.

58. Hayashi D, Shirai T, Terauchi R, Tsuchida S, Mizoshiri N, Mori Y, Arai Y, Mazda O, Kubo T. Pristimerin inhibits the proliferation of HT1080 fibrosarcoma cells by inducing apoptosis. Oncology Letters.

59. Saraswati, S.; Kumar, S.; Alhaider, A., A. α-santalol inhibits the angiogenesis and growth of human prostate tumor growth by targeting vascular endothelial growth factor receptor 2-mediated AKT/mTOR/P70S6K signaling pathway. Mol. Cancer 2013, 12(1), 147.

60. Lai, L.; Liu, J.; Zhai, D.; Lin, Q.; He, L.; Dong, Y.; Zhang, J.; Lu, B.; Chen, Y.; Yi, Z.; Liu, M. Plumbagin inhibits tumour angiogenesis and tumour growth through the Ras signalling pathway following activation of the VEGF receptor-2. Brit. J. Pharmaco. 2012, 165(4b), 1084–1096.

61. Jung, M., H.; Lee, S., H.; Ahn, E., M.; Lee, Y., M. Decursin and decursinol angelate inhibit VEGF-induced angiogenesis via suppression of the VEGFR-2-signaling pathway. Carcinogenesis 2009, 30(4), 655–661.

62. Jung, S., Y.; Choi, J., H.; Kwon, S., M.; Masuda, H.; Asahara, T.; Lee, Y., M. Decursin inhibits vasculogenesis in early tumor progression by suppression of endothelial progenitor cell differentiation and function. J. Cell. Biochem. 2012, 113(5), 1478–1487.

63. Xia Y, Min KH, Lee K. Synthesis and biological evaluation of decursin, prantschimgin and their derivatives. Bulletin of the Korean Chemical Society. 2009;30(1):43–8.

64. Lee, S.; Lee, Y., S.; Jung, S., H.; Shin, K., H.; Kim, B., K.; Kang, S., S. Anti-tumor activities of decursinol angelate and decursin fromAngelica gigas. Arch. Pharm. Res. 2003, 26(9), 727–730.

65. Pratheeshkumar, P.; Budhraja, A.; Son, Y., O.; Wang, X.; Zhang, Z.; Ding, S.; Wang, L.; Hitron, A.; Lee, J., C.; Xu, M.; Chen, G. Quercetin inhibits angiogenesis mediated human prostate tumor growth by targeting VEGFR-2 regulated AKT/mTOR/P70S6K signaling pathways. PloS ONE 2012, 7(10), e47516.

66. Miean, K., H.; Mohamed, S. Flavonoid (myricetin, quercetin, kaempferol, luteolin, and apigenin) content of edible tropical plants. J. Agr. Food Chem. 2001, 49(6), 3106–3112.

67. Le Son H, Anh NP. Phytochemical composition, in vitro antioxidant and anticancer activities of quercetin from methanol extract of Asparagus Cochinchinensis (Lour.) Merr. Tuber. Journal of Medicinal Plants Research. 2013;7(46):3360–6.

68. Nevins, J., R.; Leone, G.; DeGregori, J.; Jakoi, L. Role of the Rb/E2F pathway in cell growth control. J. Cell. Physiol. 1997, 173(2), 233–236.

69. De-Bondt, H., L.; Rosenblatt, J.; Jancarik, J.; Jones, H., D.; Morgant, D., O.; Kim, S., H. Crystal structure of cyclin-dependent kinase 2. Nature 1993, 363(6430), 595.

70. Sherr, C., J.; Roberts, J., M. CDK inhibitors: positive and negative regulators of G1-phase progression. Gene. Dev. 1999, 13(12), 1501–1512.

71. Ho, A.; Dowdy, S., F. Regulation of G1 cell-cycle progression by oncogenes and tumor suppressor genes. Curr. Opin. Genet. Dev. 2002, 12(1), 47–52.

72. Shapiro, G., I. Cyclin-dependent kinase pathways as targets for cancer treatment. J. Clin. Oncol. 2006, 24(11), 1770–1783.

73. Boonstra, J. Progression through the G1-phase of the on-going cell cycle. J. Cell. Biochem. 2003, 90(2), 244–252.

74. Harper, J., W.; Elledge, S., J. Cdk inhibitors in development and cancer. Curr. Opin. Genet. Dev. 1996, 6(1), 56–64.

75. Ullah, A.; Prottoy, N., I.; Araf, Y.; Hossain, S.; Sarkar, B.; Saha, A. Molecular Docking and Pharmacological Property Analysis of Phytochemicals from Clitoria ternatea as Potent Inhibitors of Cell Cycle Checkpoint Proteins in the Cyclin/CDK Pathway in Cancer Cells. Comput. Mol. Biosci. 2019, 9(03), 81.

76. Benson, C.; Kaye, S.; Workman, P.; Garrett, M.; Walton, M.; De-Bono, J. Clinical anticancer drug development: targeting the cyclin-dependent kinases. Br. J. Cancer 2005, 92(1), 7.

77. Deweese, J., E.; Osheroff, N. The DNA cleavage reaction of topoisomerase II: wolf in sheep’s clothing. Nucleic Acids Res. 2008, 37(3), 738–748.

78. Nitiss, J., L. Targeting DNA topoisomerase II in cancer chemotherapy. Nat. Rev. Cancer 2009, 9(5), 338.

79. Russo, P.; Del-Bufalo, A.; Cesario, A. Flavonoids acting on DNA topoisomerases: recent advances and future perspectives in cancer therapy. Curr. Med. Chem. 2012, 19(31), 5287–5293.

80. Ashour, M., E.; Atteya, R.; El-Khamisy, S., F. Topoisomerase-mediated chromosomal break repair: an emerging player in many games. Nat. Rev. Cancer 2015, 15(3), 137–151.

81. Christmann-Franck, S.; Bertrand, H., O.; Goupil-Lamy, A.; Der-Garabedian, P., A.; Mauffret, O.; Hoffmann, R.; Fermandjian, S. Structure-based virtual screening: an application to human topoisomerase II α. J. Med. Chem. 2004, 47(27), 6840–6853.

82. Folkman, J. 1984. Angiogenesis. In Biology of endothelial cells (pp. 412–428). Springer, Boston, MA.

83. Ferrara, N.; Davis-Smyth, T. The biology of vascular endothelial growth factor. Endocr. Rev. 1997, 18(1),.4–25.

84. Nowak, D., G.; Woolard, J.; Amin, E., M.; Konopatskaya, O.; Saleem, M., A.; Churchill, A., J.; Ladomery, M., R.; Harper, S., J.; Bates, D., O. Expression of pro-and anti-angiogenic isoforms of VEGF is differentially regulated by splicing and growth factors. J. Cell Sci. 2008, 121(20), 3487–3495.

85. Kim, K., J.; Li, B.; Winer, J.; Armanini, M.; Gillett, N.; Phillips, H., S.; Ferrara, N. Inhibition of vascular endothelial growth factor-induced angiogenesis suppresses tumour growth in vivo. Nature 1993, 362(6423), 841.

86. Ma, X.; Ma, C., X.; Wang, J. Endometrial carcinogenesis and molecular signaling pathways. Am. J. Mol. Biol. 2014, 4(03), 134.

87. Rini, B., I.; Vascular endothelial growth factor–targeted therapy in renal cell carcinoma: current status and future directions. Clin. Cancer Res. 2007, 13(4), 1098–1106.

88. Melincovici, C., S.; Bosca, A., B.; Susman, S.; Marginean, M.; Mihu, C.; Istrate, M.; Moldovan, I., M.; Roman, A., L.; Mihu, C., M. Vascular endothelial growth factor (VEGF)-key factor in normal and pathological angiogenesis. Rom. J. Morphol. Embryol. 2018, 59(2), 455–467.

89. Tahara M, Kiyota N, Yamazaki T, Chayahara N, Nakano K, Inagaki L, Toda K, Enokida T, Minami H, Imamura Y, Sasaki T. Lenvatinib for anaplastic thyroid cancer. Frontiers in oncology. 2017 Mar 1;7:25.

90. Roskoski-Jr, R.; Vascular endothelial growth factor (VEGF) and VEGF receptor inhibitors in the treatment of renal cell carcinomas. Pharmacol. Res. 2017, 120, 116–132.

91. Schrödinger Release 2018-4: Protein Preparation Wizard; Epik, Schrödinger, LLC, New York, NY, 2016; Impact, Schrödinger, LLC, New York, NY, 2016; Prime, Schrödinger, LLC, New York, NY, 2018.

92. Schrödinger Release 2018-4: Prime, Schrödinger, LLC, New York, NY, 2018.

93. Schrödinger Release 2018-4: LigPrep, Schrödinger, LLC, New York, NY, 2018.

94. Schrödinger Release 2018-4: Epik, Schrödinger, LLC, New York, NY, 2018.

95. Schrödinger Release 2018-4: Glide, Schrödinger, LLC, New York, NY, 2018.

96. Dash R, Hosen SZ, Karim MR, Kabir MS, Hossain MM, Junaid M, Islam A, Paul A, Khan MA. In silico analysis of indole-3-carbinol and its metabolite DIM as EGFR tyrosine kinase inhibitors in platinum resistant ovarian cancer vis a vis ADME/T property analysis. J App Pharm Sci. 2015 Nov;5(11):073–8.

97. Visualizer, D.S. (2017) Release 4.1. Accelrys Inc., San Diego, CA.

98. Daina A, Michielin O, Zoete V. SwissADME: a free web tool to evaluate pharmacokinetics, drug-likeness and medicinal chemistry friendliness of small molecules. Scientific reports. 2017 Mar 3;7:42717.

99. Cheng, F.; Li, W.; Zhou, Y.; Shen, J.; Wu, Z.; Liu, G.; Lee, P., W.; Tang, Y. 2012. admetSAR: a comprehensive source and free tool for assessment of chemical ADMET properties.

100. Dong, J.; Wang, N., N.; Yao, Z., J.; Zhang, L.; Cheng, Y.; Ouyang, D.; Lu, A., P.; Cao, D., S. ADMETlab: a platform for systematic ADMET evaluation based on a comprehensively collected ADMET database. J. Cheminformatics 2018, 10(1), 29.

101. Filimonov, D., A.; Lagunin, A., A.; Gloriozova, T., A.; Rudik, A., V.; Druzhilovskii, D., S.; Pogodin, P., V.; Poroikov, V., V. Prediction of the biological activity spectra of organic compounds using the PASS online web resource. Chem. Heterocycl. Compd. 2014, 50(3), 444–457.

102. Geronikaki, A.; Poroikov, V.; Hadjipavlou-Litina, D.; Filimonov, D.; Lagunin, A.; Mgonzo, R. Computer aided predicting the biological activity spectra and experimental testing of new thiazole derivatives. Quant. Struct.-act. Rel. 1999, 18(1), 16–25.

103. Zaretzki, J.; Bergeron, C.; Huang, T., W.; Rydberg, P.; Swamidass, S., J.; Breneman, C., M. RS-WebPredictor: a server for predicting CYP-mediated sites of metabolism on drug-like molecules. Bioinformatics 2012, 29(4), 497–498.

104. Drwal, M., N.; Banerjee, P.; Dunkel, M.; Wettig, M., R.; Preissner, R. ProTox: a web server for the in silico prediction of rodent oral toxicity. Nucleic Acids Res. 2014, 42(W1), W53–W58.

105. Schrödinger Release 2018-4: Jaguar, Schrödinger, LLC, New York, NY, 2018.

106. Lee, C.; Yang, W.; Parr, R., G. Development of the Colle-Salvetti correlation-energy formula into a functional of the electron density. Phys. Rev. B 1988, 37(2), 785.

107. Becke, A., D. Density-functional exchange-energy approximation with correct asymptotic behavior. Phys. Rev. A 1988, 38(6), 3098.

108. Pearson, R., G. Absolute electronegativity and hardness correlated with molecular orbital theory. P. Natl. Acad. Sci. 1986, 83(22), 8440–8441.

109. Parr, R., G.; Yang, W. Density-Functional Theory of Atoms and Molecules, vol. 16 of International series of monographs on chemistry. Oxford University Press 1989, New York.

110. file:///C:/Program%20Files/Schrodinger20184/docs/Documentation.htm#maestro_tools_help/ramachandran_panel.html

111. Yuriev, E.; Ramsland, P., A. Latest developments in molecular docking: 2010–2011 in review. J. Mol. Recognit. 2013, 26(5), 215–239.

112. Zhang, X.; Perez-Sanchez, H.; C-Lightstone, F. A comprehensive docking and MM/GBSA rescoring study of ligand recognition upon binding antithrombin. Curr. Top. Med. Chem. 2017, 17(14), 1631–1639.

113. Sherman, W.; Day, T.; Jacobson, M., P.; Friesner, R., A.; Farid, R. Novel procedure for modeling ligand/receptor induced fit effects. J. Med. Chem. 2006, 49(2), 534–553.

114. Aamir, M.; Singh, V., K.; Dubey, M., K.; Meena, M.; Kashyap, S., P.; Katari, S., K.; Upadhyay, R., S.; Singh, S. In silico Prediction, Characterization, Molecular Docking and Dynamic Studies on Fungal SDRs as Novel Targets for Searching Potential Fungicides against Fusarium Wilt in Tomato. Front. Pharmacol. 2018, 9, 1038.

115. Friesner, R., A.; Murphy, R., B.; Repasky, M., P.; Frye, L., L.; Greenwood, J., R.; Halgren, T., A.; Sanschagrin, P., C.; Mainz, D., T. Extra precision glide: Docking and scoring incorporating a model of hydrophobic enclosure for protein− ligand complexes. J. Med. Chem. 2006, 49(21), 6177–6196.

116. Priyadarshini, V.; Pradhan, D.; Munikumar, M.; Swargam, S.; Umamaheswari, A.; Rajasekhar, D. Genome-based approaches to develop epitope-driven subunit vaccines against pathogens of infective endocarditis. J. Biomol. Struct. Dyn. 2014, 32(6), 876–889.

117. Lipinski, C., A.; Lombardo, F.; Dominy, B., W.; Feeney, P., J. Experimental and computational approaches to estimate solubility and permeability in drug discovery and development settings. Adv. Drug Deliv. Rev. 1997, 23(1-3), 3–25.

118. Gohlke H, Hendlich M, Klebe G. Knowledge-based scoring function to predict protein-ligand interactions. Journal of molecular biology. 2000 Jan 14;295(2):337–56.

119. Shoichet BK, McGovern SL, Wei B, Irwin JJ. Lead discovery using molecular docking. Current opinion in chemical biology. 2002 Aug 1;6(4):439–46.

120. Klebe G. Protein-ligand interactions as the basis for drug action. InMultifaceted Roles of Crystallography in Modern Drug Discovery 2015 (pp. 83–92). Springer, Dordrecht.

121. Lipinski, C., A.; Lead-and drug-like compounds: the rule-of-five revolution. Drug Discov. S Today Technol. 2004, 1(4), 337–341.

122. Pollastri, M., P. Overview on the Rule of Five. Curr. Protoc. Pharmacol. 2010, 49(1), 9–12.

123. Li, A., P. Screening for human ADME/Tox drug properties in drug discovery. Drug Discov. Today 2001, 6(7), 357–366.

124. Guengerich, F., P. Cytochrome P-450 3A4: regulation and role in drug metabolism. Annu. Rev. Pharmacol. Toxicol. 1999, 39(1), 1–17.

125. Glue, P.; Clement, R., P. Cytochrome P450 enzymes and drug metabolism—basic concepts and methods of assessment. Cell. Mol. Neurobiol. 1999, 19(3), 309–323.

126. Dixit, B. A review on the effects of CMPF binding with Human Serum Albumin. Bioinformatics Rev. 2017, 3(9), 9–18.

127. Radchenko, E., V.; Dyabina, A., S.; Palyulin, V., A.; Zefirov, N., S. Prediction of human intestinal absorption of drug compounds. Russ. Chem. Bull. 2016, 65(2), 576–580.

128. Wessel, M., D.; Jurs, P., C.; Tolan, J., W.; Muskal, S., M. Prediction of human intestinal absorption of drug compounds from molecular structure. J. Chem. Inf. Comput. Sci. 1998, 38(4), 726–735.

129. Basant, N.; Gupta, S.; Singh, K., P. Predicting human intestinal absorption of diverse chemicals using ensemble learning based QSAR modeling approaches. Comput. Biol. Chem. 2016, 61, 178–196.

130. Swierczewska, M.; Lee, K., C.; Lee, S. What is the future of PEGylated therapies?. 2015

131. Smalling, R., W. Molecular biology of plasminogen activators: what are the clinical implications of drug design?. Am. J. Cardiol. 1996, 78(12), 2–7.

132. Sahin, S.; Benet, L., Z. The operational multiple dosing half-life: a key to defining drug accumulation in patients and to designing extended release dosage forms. Pharm. Res. 2008, 25(12), 2869–2877.

133. Sanguinetti, M., C.; Jiang, C.; Curran, M., E.; Keating, M., T. A mechanistic link between an inherited and an acquird cardiac arrthytmia: HERG encodes the IKr potassium channel. Cell 1995, 81(2), 299–307.

134. Aronov, A., M. Predictive in silico modeling for hERG channel blockers. Drug Discov. Today 2005, 10(2), 149–155.

135. Cheng, A.; Dixon, S., L. In silico models for the prediction of dose-dependent human hepatotoxicity. J. Comput. Aid. Mol. Des. 2003, 17(12), 811–823.

136. Xu, J., J.; Diaz, D.; O’Brien, P., J. Applications of cytotoxicity assays and pre-lethal mechanistic assays for assessment of human hepatotoxicity potential. Chem. Biol. Interact. 2004, 150(1), 115–128.

137. Mortelmans, K.; Zeiger, E. The Ames Salmonella/microsome mutagenicity assay. Mutat. Res-Fund Mol. M. 2000, 455(1-2), 29–60.

138. Holt, M., P.; Ju, C. Mechanisms of drug-induced liver injury. AAPS J. 2006, 8(1), E48–E54.

139. Lagunin, A.; Stepanchikova, A.; Filimonov, D.; Poroikov, V. PASS: prediction of activity spectra for biologically active substances. Bioinformatics 2000, 16(8), 747–748.

140. United Nations. Economic Commission for Europe. Secretariat, 2005. Globally harmonized system of classification and labelling of chemicals (GHS). United Nations Publications.

141. Tyzack, J., D.; Mussa, H., Y.; Williamson, M., J.; Kirchmair, J.; Glen, R., C. Cytochrome P450 site of metabolism prediction from 2D topological fingerprints using GPU accelerated probabilistic classifiers. J. scinemrofnimehC 2014, 6(1), 29.

142. Danielson, P., B. The cytochrome P450 superfamily: biochemistry, evolution and drug metabolism in humans. Curr. Drug Metab. 2002, 3(6), 561–597.

143. Matysiak, J. Evaluation of electronic, lipophilic and membrane affinity effects on antiproliferative activity of 5-substituted-2-(2, 4-dihydroxyphenyl)-1, 3, 4-thiadiazoles against various human cancer cells. Eur. J. Med. Chem. 2007, 42(7), 940–947.

144. Zhan, C., G.; Nichols, J., A.; Dixon, D., A. Ionization potential, electron affinity, electronegativity, hardness, and electron excitation energy: molecular properties from density functional theory orbital energies. J. Phys. Chem. A 2003, 107(20), 4184–4195.

145. Hoque, M., M.; Halim, M., A.; Sarwar, M., G.; Khan, M., W. Palladium-catalyzed cyclization of 2-alkynyl-N-ethanoyl anilines to indoles: synthesis, structural, spectroscopic, and mechanistic study. J. Phys. Org. Chem. 2015, 28(12), 732–742.

146. Ayers, P., W.; Parr, R., G.; Pearson, R., G. Elucidating the hard/soft acid/base principle: a perspective based on half-reactions. J. Chem. Phys. 2006, 124(19), 194107.

147. Sarkar B, Islam SS, Ullah MA, Hossain S, Prottoy MN, Araf Y, Taniya MA. Computational Assessment and Pharmacological Property Breakdown of Eight Patented and Candidate Drugs against Four Intended Targets in Alzheimer’s Disease. Advances in Bioscience and Biotechnology. 2019 Nov 25;10(11):405. DOI: https://doi.org/10.4236/abb.2019.1011030

148. Ullah MA, Johora FT, Sarkar B, Araf Y, Rahman MH. Curcumin Analogues as the Inhibitors of TLR4 Pathway in Inflammation and Their Drug Like Potentialities: A Computer-based Study. BioRxiv. 2020 Jan 1. DOI: https://doi.org/10.1101/2020.01.27.921528

